# Transcription factors protect from DNA re-methylation during reprograming of primordial germ cells and pre-implantation embryos

**DOI:** 10.1101/850362

**Authors:** Isaac Kremsky, Victor G. Corces

**Author notes:** Corresponding Author: Victor G. Corces.

## Abstract

A growing body of evidence suggests that certain phenotypic traits of epigenetic origin can be passed across generations via both the male and female germlines of mammals. These observations have been difficult to explain owing to a global loss of the majority of known epigenetic marks present in parental chromosomes during primordial germ cell development and after fertilization. By integrating previously published BS-seq, DNase-seq, ATAC-seq, and RNA-seq data collected during multiple stages of primordial germ cell and preimplantation development, we find that the methylation status of the majority of CpGs genome-wide is restored after global reprogramming, despite the fact that global CpG methylation drops to 10% in primordial germ cells and 20% in the inner cell mass of the blastocyst. We estimate the proportion of such CpGs with preserved methylation status to be 78%. Further, we find that CpGs at sites bound by transcription factors during the global re-methylation phases of germ line and embryonic development remain hypomethylated across all developmental stages observed. On the other hand, CpGs at sites not bound by transcription factors during the global re-methylation phase have high methylation levels prior to global de-methylation, become de-methylated during global de-methylation, and then become re-methylated. The results suggest that transcription factors can act as carriers of epigenetic information during germ cell and pre-implantation development by ensuring that the methylation status of CpGs is maintained after reprogramming of DNA methylation. Based on our findings, we propose a model in which transcription factor binding during the re-methylation phases of primordial germ cell and pre-implantation development allow epigenetic information to be maintained trans-generationally even at sites where DNA methylation is lost during global de-methylation.

## Background

Evidence for the transmission of phenotypic traits via inter- and trans-generational epigenetic inheritance in mammals has grown substantially in recent years. However, the underlying mechanisms responsible for these phenomena have remained elusive [1, 2]. DNA methylation is arguably the best candidate carrier of epigenetic information and an appealing option to explain inter- and trans-generational epigenetic inheritance, since it is heritable across rounds of DNA replication. However, the genome is globally de-methylated in mammals at two developmental stages – the primordial germ cell (PGC) and the pre-implantation embryo stages. Global DNA methylation levels fall below 10% in PGCs [3], and below 20% in the inner cell mass (ICM) stage of pre-implantation embryos [4]. Maintenance of CpG methylation during PGC and preimplantation development at endogenous retroviral elements of the Intracisternal A Particle (IAP) type are known to be involved in some cases of tans-generationally inherited epialleles, the agouti viable yellow and axin fused loci [5–8]. In these examples, the phenotype is correlated with the DNA methylation status of the IAP inserted near the relevant genes, driving their expression via an alternate promoter within the IAP long terminal repeat (LTR) [7, 9]. CpGs at IAPs are known to be resistant to global de-methylation during PGC development [3], thus maintenance of CpG methylation status at these loci across PGC and pre-implantation development is the likely mechanism of epigenetic inheritance in these examples.

While it is possible that the remaining 10% of methylated CpGs during PGC development is sufficient to explain inter- and trans-generational epigenetic inheritance of all known heritable epiphenotypes, genome-wide BS-seq studies show that the methylation status of a substantial proportion of CpG sites is faithfully recapitulated before and after global de-methylation and re-methylation across both PGC [3] and embryonic [4] development. That is, a substantial number of CpGs that are methylated prior to global de-methylation are first de-methylated and then re-methylated, whereas many CpGs that are not methylated prior to global de-methylation remain unmethylated even after global re-methylation. This suggests that additional carrier(s) of epigenetic information must exist to maintain the memory of CpG methylated states after reprogramming events. Identifying the additional carrier(s) is essential for understanding mechanisms of inter- and trans-generational epigenetic inheritance. The extent to which CpG methylation status is faithfully maintained has not been quantified before.

A number of candidate carriers of epigenetic information besides DNA methylation have been proposed to be involved in mechanisms of inter- and trans-generational epigenetic inheritance, including miRNAs, piRNAs, and tRNAs [2, 10–13]. However, mechanisms explaining these phenomena are generally thought to require that epigenetic information is passed between different types of molecular carriers over the course of development [14]. Although alterations in sperm non-coding RNAs (ncRNAs) have been shown to induce trans-generational effects on phenotypes [10, 11, 13], only a few studies have observed alterations in sperm RNAs beyond the F_1_ generation. For example, mice subjected to stress have altered miRNAs in F_1_, but not F_2_ sperm, despite the fact that F_3_ mice still displayed the same alterations in phenotype that were observed in the F_1_ and F_2_ generations due to stress in the F_0_ [10]. This suggests that alterations in miRNAs are capable of inducing a trans-generational effect on phenotype, but that some other molecular carrier may be responsible for the trans-generational maintenance of the altered phenotype [10].

Histone modifications are widely studied molecular carriers of epigenetic information.Studies have found that H3K4me3 and H3K27me3 can be passed on from oocytes to pre-implantation embryos, but the same studies found that these marks are globally reprogrammed on paternal alleles upon fertilization [15, 16]. To date no studies have identified a histone modification present in sperm that is maintained upon fertilization. Therefore, while histone modifications could act in mechanisms of maternal inter- and trans-generational epigenetic inheritance, they may not sufficient to explain the transmission of altered epigenetic states through the paternal gametes. Molecular carriers other than CpG methylation involved in trans-generational maintenance of epi-alleles have to this point still not been identified. Nucleosomes containing H2A.Z and H3K4me1 have been shown to antagonize de-novo CpG methylation in zebrafish during pre-implantation development [17], and it is possible that a similar mechanism exists during the re-methylation phases in mammals.

Transcription factors (TFs) could also serve as carriers of epigenetic information but their possible role in this process has received only moderate attention. TFs are present on the genomes of mature gametes [18, 19], and there is evidence that TF binding can influence DNA methylation at its binding site, both by direct steric hindrance of DNA methyltransferases and by recruitment of Tet enzymes to specific TFs bound to DNA [20]. In addition, a class of TFs known as pioneer factors can bind to nucleosomes and can stabilize nucleosome positioning [21].Therefore, TFs could also be involved in the placement of marker nucleosomes to inhibit de novo DNA methylation. TFs thus represent a plausible candidate that could pass on epigenetic information when DNA methylation is erased, acting as mediators that can allow cells to preserve DNA methylation patterns across PGC differentiation and pre-implantation development.

In order to test the hypothesis that TFs might act as mediators of epigenetic memory during DNA methylation reprogramming we integrated publicly available BS-seq [3, 4], RNA-seq [3, 15], DNase-seq [22, 23], and ATAC-seq [19, 24, 25] data across multiple stages of PGC and embryonic development. We find a striking correlation between the presence or absence of bound TFs during global re-methylation of PGCs and ESCs and the methylation status of CpGs at their core binding sequence, both before and after embryonic development. The results suggest that TFs may be involved in the maintenance of epigenetic information during global hypomethylation of PGCs and ESCs, ensuring that the methylation state of the majority of CpGs is preserved between generations across embryonic development. This TF-mediation model of epigenetic inheritance leads to testable predictions for how both inter- and trans-generational epigenetic inheritance might occur mechanistically.

## Results

### TF binding at CpGs in E14.5 male PGCs predicts DNA methylation patterns throughout male PGC development

PGCs first appear in the epiblast at E6.5 of mouse embryonic development and, in male embryos, they eventually become sperm. As PGCs migrate to the genital ridge and differentiate, their genomes become de-methylated, and de-methylation is completed by E13.5, with 10% of CpGs in the genome still remaining methylated [3], mostly at transposable element sequences. After this time, male PGCs become remethylated, and average levels of methylation reach 50% by E16.5 [3] and nearly 80% in the mature sperm [4]. The paternal genome is largely demethylated immediately after fertilization whereas the maternal genome is demethylated more slowly during pre-implantation development. Both genomes retain an average of 20% of CpG methylation in the ICM of the blastocyst at E3.5 [4]. Re-methylation of the paternal and maternal chromosomes takes place during subsequent embryonic development and ~70% of CpGs are methylated in the epiblast at E6.5 [3]. We sought to quantify the degree to which the methylation status of DNA is preserved across PGC and preimplantation development by comparing the CpG methylation state of E6.5 epiblast cells with that of sperm, with the caveats that only a subset of epiblast cells give rise to the germline and that epiblast cells have not yet undergone complete re-methylation. Using previously published genome-wide BS-seq data (GWBS) [3, 4], we find that 91% of CpGs with very high (>80%) methylation in the epiblast also have the same methylation level in sperm, whereas 90% of CpGs that have very high methylation in sperm have >50% methylation in epiblast cells (Figure 1A). Similarly, of the CpGs with very low (< 20%) methylation in epiblasts, 83% also have very low methylation in sperm, whereas 47% of CpGs with very low methylation in sperm also have very low methylation in epiblasts (Figure 1A). Given that epiblasts are still undergoing global de-novo methylation whereas sperm are fully methylated, the methylation levels in sperm are more reliable to estimate the proportion of CpGs whose methylation status is faithfully maintained after reprogramming. Although BS-seq data in successive generations of sperm are needed to precisely determine the amount of CpGs with preserved methylation status across generations, comparing the epiblast data to sperm can at least give us a sense and reasonable estimate of the amount of preserved CpGs. Of the high confidence CpGs in sperm, only 9% had a methylation level in between 80% and 20%, i.e. roughly 91% of CpGs in sperm have either > 80% or less than 20% methylation. Conservatively assuming that the 9% of intermediately methylated CpGs in sperm are not preserved across generations, we estimate (see Methods) that the methylation level of 78% of CpGs is preserved before and after the global de-methylation and re-methylation phases of PGC and preimplantation development and, therefore, across generations.

**Fig. 1.**
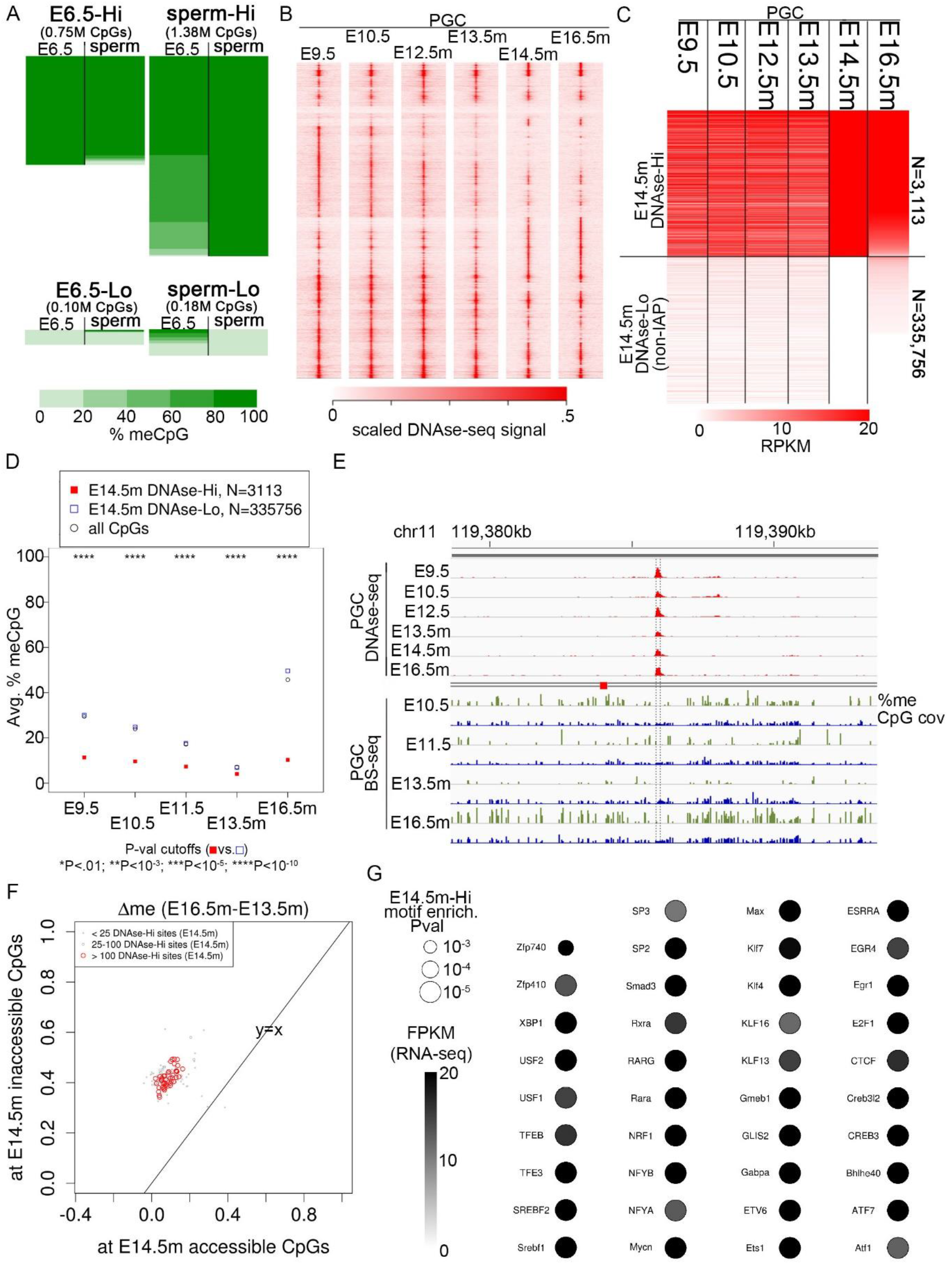
TF binding at CpGs in E14.5m PGCs predicts DNA methylation patterns throughout male PGC development**. A** Heatmaps of % methylation from BS-seq signal in E6.5 epiblasts and sperm. Each row represents a CpG with at least 10 BS-seq reads in both samples. The heatmaps are divided as follows: CpGs with very high methylation (> 80%) in epiblast sorted by % methylation in sperm (top left); CpGs with very low methylation (< 20%) in epiblasts, sorted by methylation in sperm (bottom left); CpGs with very high methylation in sperm, sorted by methylation in epiblasts (top right); CpGs with very low methylation in sperm, sorted by methylation in epiblasts (bottom right). The length of each heatmap is scaled to the total number of CpGs within it. **B** Heatmap of DNase-seq signal centered at binding sequences called by fimo that overlap a called peak summit in at least one of the samples displayed, clustered by hierarchical clustering. **C** Heatmap of RPKM values within just binding sequences called by fimo, separated into those that have high DNase-signal in E14.5m (DNase-Hi, RPKM > 20) and those that have RPKM=0 (DNase-Lo). Rows are ordered by decreasing signal in E16.5m. **D** Average methylation of CpGs within E14.5m DNase-Hi and DNase-Lo regions, weighted by the number of BS-seq reads at each CpG. P values are by Fisher’s exact test. The numbers displayed correspond to the number of TF binding sequences at which CpG methylation was averaged. **E** Example region with DNase accessibility and low local DNA methylation throughout PGC development. CpG cov indicates the BS-seq read coverage. **F** Scatterplot of changes in DNA methylation between E14.5m and E16.5m PGCs. Each point represents the average methylation change (fraction meCpG in E16.5m minus E13.5m PGCs) at CpGs overlapping a binding sequence for a specific TF. The x-axis gives the average value for E14.5m PGC DNase-Hi sites, while the y-axis gives the average value for DNase-Lo sites. **G** Significantly enriched motifs at E14.5m DNase-Hi sites. P-values are by Fisher’s Exact test.

The mechanisms by which PGCs retain a memory of the previous DNA methylation state after global DNA de-methylation are unclear. We thus sought to test the hypothesis that the presence of DNA-bound TFs precludes re-methylation of bound sequences. To this end, we used publicly available DNase-seq data obtained in PGCs at different stages of differentiation, from days of embryonic development E9.5-E13.5, when DNA de-methylation is completed, to E14.5-E16.5 when re-methylation is almost finished in male PGCs [22]. We analyzed the pattern of TF dynamics at distal CpG sites (>2.5 kb from annotated TSSs at sites presumed to be enhancers) using unsupervised clustering and found a number of distinct temporal patterns of TF occupancy (Figure 1B). Some TF binding sites remain occupied throughout PGC development, whereas others are either gained or lost at specific stages. However, the TF binding status remains largely fixed between embryonic days E14.5 and E16.5 of male PGC development (E14.5m and E16.5m, respectively) (Figure 1B). To determine the relationship between TF occupancy and DNA methylation, we selected TF binding sequences with high DNase-seq signal in E14.5m PGCs (DNase-Hi), and compared them to a control set of TF binding sites with no DNase-seq signal in E14.5m (DNase-Lo) (Figure 1C). To identify DNase-Lo sites, we obtained DNase-seq peaks for all available mouse tissues from ENCODE [26], including all peaks from the PSU Hardison [27] and UW Stamatoyannopoulous [28–30] labs. We merged all ENCODE peaks with all DNase-seq peaks from PGCs and ESCs, and we used fimo [31] to identify DNase-seq peaks containing known TF binding sequences. DNase-Hi and -Lo sites consist only of the merged set of binding sequences themselves, and not the rest of the peak regions, since those are the presumed binding sites of the TFs. The level of DNase-seq signal at DNase-Hi and -Lo sites is largely unchanged between E14.5m and E16.5m but is distinct at all other PGC stages (Figure 1C; Additional file 1: Figure S1A), suggesting that TFs remain persistently bound during the re-methylation phase of PGC development.

We then compared the average DNA methylation levels at DNase-Hi and DNase-Lo CpGs across PGC development (Figure 1D,E; Additional file 1: Figure S1B,C). The DNA methylation pattern at the DNase-Lo sites resembles that of the global average: they are progressively de-methylated from E9.5 to E13.5m, and then re-methylated thereafter. On the other hand, DNase-Hi sites remain largely unmethylated at all stages of PGC development for which data exist, and methylation levels at the DNase-Hi sites are significantly lower than those at DNase-Lo sites at all PGC stages. We then examined the average change in methylation from E13.5m to E16.5m at binding site sequences for each TF separately, limiting the set of TFs to only those that have RNA-seq signal at E16.5m. For every such TF binding sequence, sites with high DNase-seq signal specifically at the binding sequence had a smaller increase in DNA methylation levels than did sites that were not bound by TFs (Figure 1F). A motif enrichment analysis revealed 39 significantly enriched TFs that are highly expressed in E16.5m PGCs (Figure 1G; Additional file 1: Figure S1D; Additional file 2: Table S1). The majority of these TFs have clear DNase footprints at their DNase-Hi sites (Additional file 3), suggesting that TFs are bound specifically at the majority of DNase-Hi sites identified in E14.5m.

Taken together, these results support the hypothesis that TFs can act as mediators of epigenetic information during PGC development by affecting DNA methylation levels. CpGs bound by a TF at E14.5m tend to be unmethylated throughout PGC development, whereas CpGs that do not bind a TF at E14.5m are methylated early in PGC development, become progressively de-methylated until E13.5, and then become remethylated in males. Females do not undergo re-methylation until much later than E13.5 and so we were unable to test this phenomenon in females.

### Global TF binding and DNA methylation patterns during male PGC and ESC development are highly correlated

The fact that approximately 78% of CpGs preserve their methylation status between sperm and E6.5 epiblasts (Figure 1A) suggests that the methylation status is preserved not just across global de-methylation followed by re-methylation of PGCs, but also across global de- and re-methylation in preimplantation embryos after fertilization. We therefore wondered if the same pattern of TF binding during re-methylation of PGCs would also be present during re-methylation of ESCs. To test this, we examined ATAC-seq signal in embryonic stem cells (ESCs) [24] at the E14.5m PGC DNase-Hi sites. We found that the vast majority of E14.5m DNase-Hi sites have either intermediate or high ATAC-seq signal in ESCs (Figure 2A, B). Less than 5% of E14.5m PGC DNase-Hi sites have low ATAC-seq signal in ESCs (Figure 2B).

**Fig. 2.**
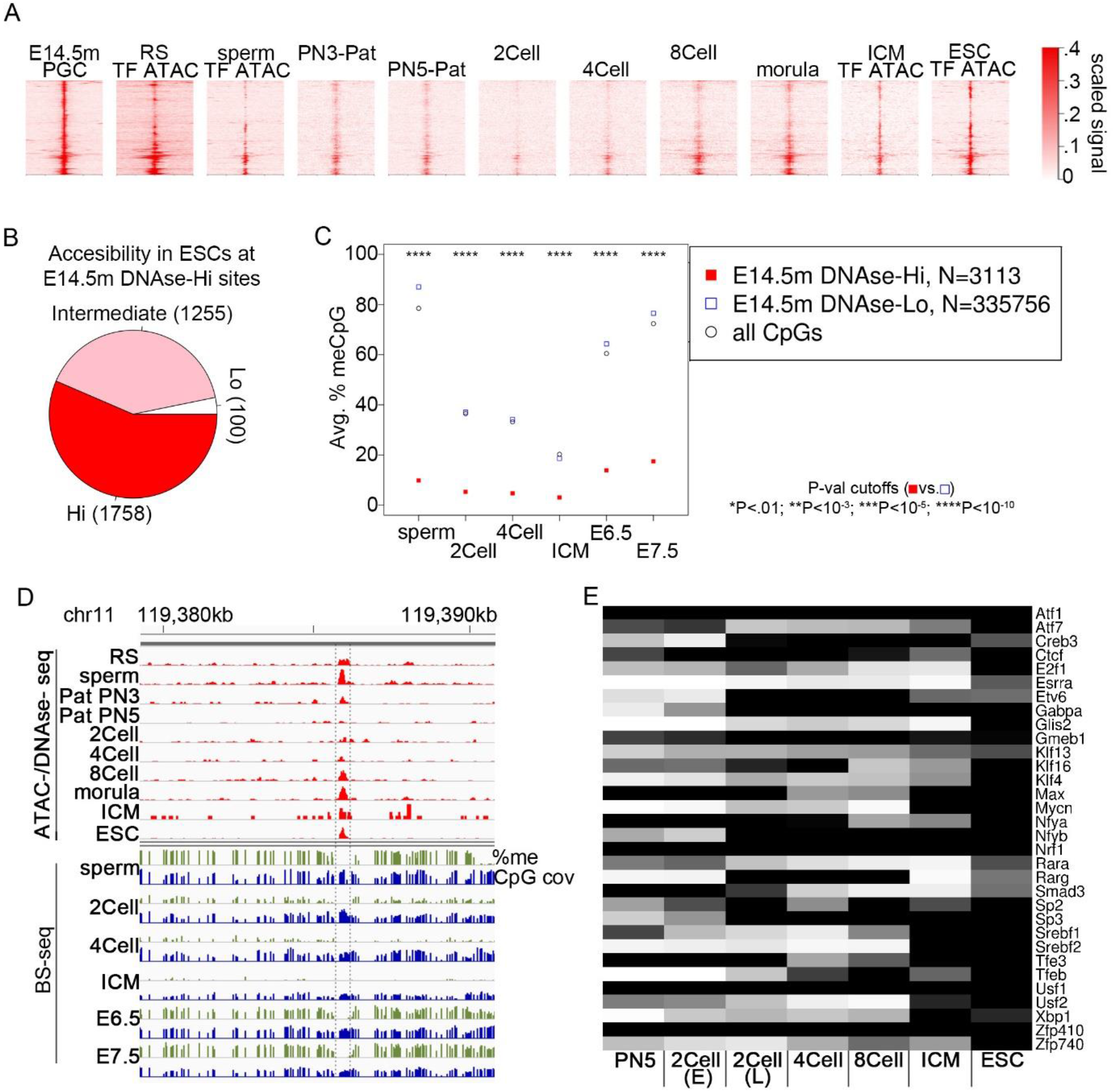
Global TF binding and CpG methylation patterns during male PGC and ESC development are highly correlated. For all cases where ATAC-seq signal was used in this figure, only ATAC-seq fragments < 115 bp, which indicate TF binding (TF-ATAC), were used. **A** Heatmap at the indicated stages of gamete and embryonic development, centered at E14.5m PGC DNase-Hi sites ordered by hierarchical clustering. RS=round spermatid. PN3_P, PN5_P=Paternal PN3 and PN5 pronucleus, respectively. Data from TF-ATAC are labelled as such; all other data are from DNAse-seq. **B** Pie chart showing the proportion of E14.5m PGC DNase-Hi sites with high, intermediate, or low ATAC-seq signal in ESCs. **C** Same as Fig. 1D except that the stages at which the average CpG methylation are calculate are at gamete and embryonic development of the indicated stages. **D** Example displaying DNase-seq and BS-seq signal at the same region as in Fig. 1D. **E** Heatmap of RNA-seq signal at the indicated stages for the significantly enriched TFs at E14.5m PGC DNase-Hi sites from Fig. 1F.

We next examined the DNA methylation levels throughout pre-implantation development at E14.5m PGC DNase-Hi and DNase-Lo sites using published BS-seq data [4]. We found the same behavior as in PGCs: the DNase-Lo sites follow the global average pattern of de-methylation after fertilization and up to the ICM stage, followed by re-methylation thereafter, whereas the DNase-Hi sites remain lowly methylated at all stages for which data exist (Figure 2C,D; Additional file 1: Figure S2A-C). Binding sequences for the same set of 39 TFs identified as putative epigenetic mediators of PGC development (Figure 1G) show globally less change in DNA methylation levels between ICMs and E7.5 embryos when bound by a TF than when unbound (Additional file 1: Figure S2A). As is the case in PGCs, most of these TFs show clear footprints at their binding sequences in ESCs (Additional file 4). Each of these 39 TFs is highly expressed in ESCs based on RNA-seq data (Figure 2E).

Taken together, these results, along with those discussed in the previous section, suggest that, despite two phases of global de-methylation followed by re-methylation, the methylation status of the majority of CpGs in the mammalian genome is preserved across embryonic development, and that the binding of TFs during the re-methylation phases is associated with maintenance of hypomethylation across development. An estimated 78% of the CpGs in the genome behave in this manner. On the other hand, unbound TF binding sites are methylated both before and after the two rounds of global de-/re-methylation.

### CpG methylation reprogramming across PGC development is associated with the binding of putative reprogramming TFs

Given that the majority of CpGs have a preserved methylation status across embryonic development, we sought to characterize the CpGs whose methylation status is not preserved. There is evidence that some TFs, including CTCF and Esrrb, can bind to methylated DNA and are associated with de-methylation of their binding sites and the surrounding regions [32–35]. We thus wondered whether this is also the case during PGC development. To test this, we took the set of regions with low DNase-seq signal in E9.5 PGCs (E9.5-trace) but high DNase-seq signal in E14.5m PGCs (E14.5m-Hi), and compared them to regions with high signal at E9.5 PGCs and low signal at E14.5m PGCs (E9.5-Hi, E14.5m-trace). Both TF binding and DNA methylation were compared between these sites throughout germline and embryonic development (Figure 3A-F). We found that the DNA methylation signal was reprogrammed according to its change in DNase-seq signal: E9.5-Hi, E14.5m-trace sites have low DNA methylation at E9.5 but high DNA methylation at E16.5m PGCs, whereas E9.5-trace, E14.5m-Hi sites have high DNA methylation levels at E9.5 and low DNA methylation levels at E16.5m (Figure 3A,B). Both E9.5-Hi, E14.5m-trace and E9.5-trace, E14.5m-Hi sites show no discernible pattern of TF binding during preimplantation development and in sperm (Figure 3C). However, the DNA methylation levels of E9.5-Hi, E14.5m-trace sites remain high in sperm, consistent with their high levels in E16.5m PGCs. They then undergo rapid de-methylation after fertilization and are only methylated at intermediate levels in the epiblast stage (Figure 3D). Since they are lowly methylated in E9.5 PGCs, this suggest the DNA methylation is reprogrammed sometime between the epiblast and early PGC stages, hinting at a possible role for these sites in early PGC development. On the other hand, E9.5-trace, E14.5m-Hi sites have low DNA methylation levels in sperm, consistent with their low levels in E16.5m PGCs, but then become highly methylated in the epiblast stage. This suggests that the E9.5-trace, E14.5m-Hi sites are reprogrammed between E13.5 and E16.5 in male PGCs (Figure 3D).

**Fig. 3.**
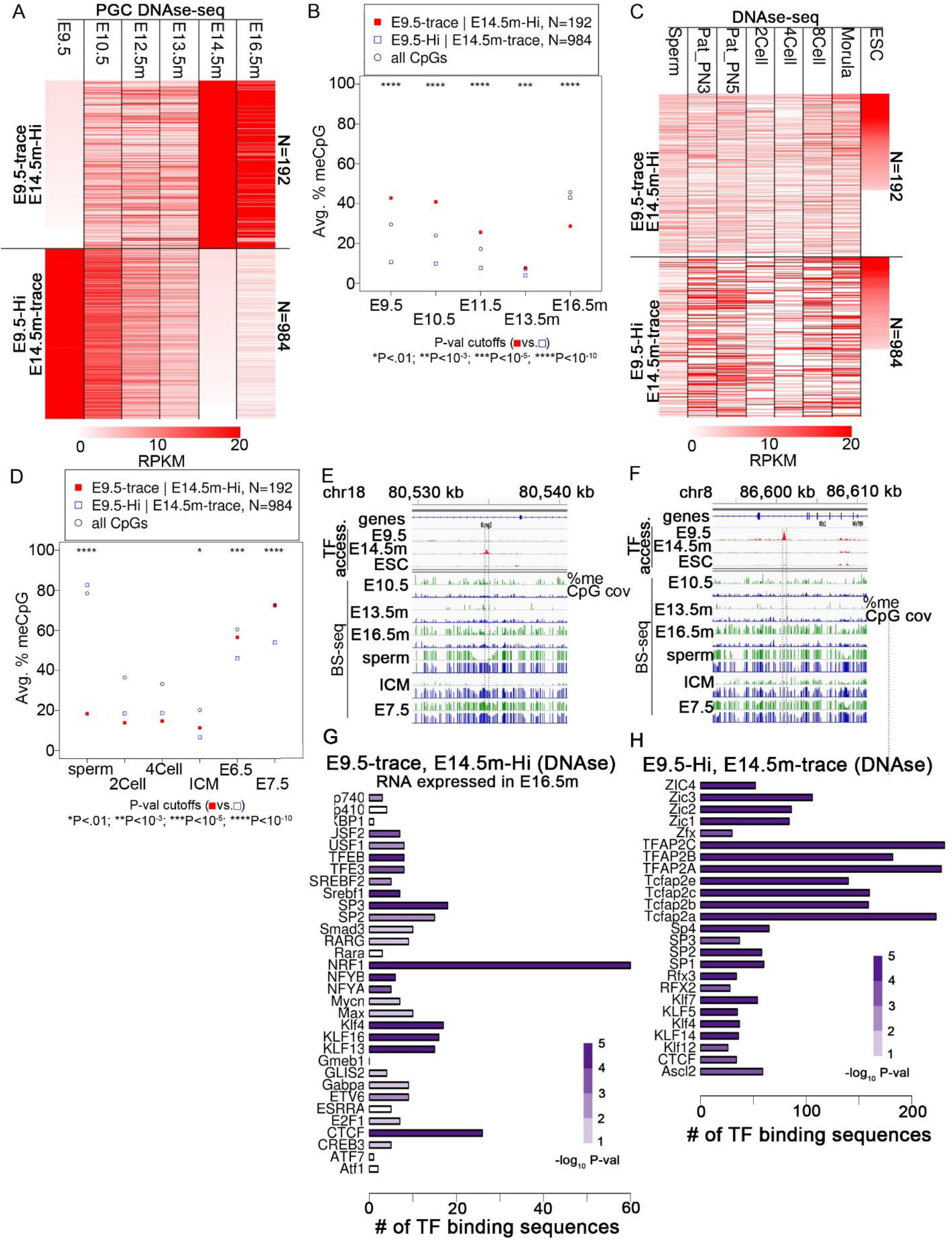
CpG methylation is reprogrammed only at a small fraction of TF binding sites, at binding sites for specific reprogramming TFs during male PGC and ESC development. **A** Heatmaps of DNase-seq signal at the indicated sites during PGC development. E9.5-trace, E14.5m-Hi sites are those that have FPKM<1 in E9.5 PGCs and FPKM>20 in E14.5E14.5m PGCs, while E9.5-Hi, E14.5m-trace sites have FPKM>20 in E9.5 PGCs and FPKM<1 in E14.5m PGCs. **B** Average DNA methylation levels at the indicated regions at the indicated stages of PGC development, as described in Fig.1D. **C** Same as A, except during gamete and preimplantation development. **D** Same as B, except during gamete and preimplantation development. **E** Example showing the DNA methylation across development at an E9.5-trace, E14.5m-Hi site with a CTCF binding sequence. **F** Example showing the DNA methylation across development at an E9.5-Hi,E14.5m-trace site with a CTCF binding sequence. CpG cov indicates the BS-seq read coverage. **G,H** Bar plots showing the number of motif hits for each TF at the indicated set of regions. Statistical significance of the enrichment of each TF motif, based on Fisher’s exact test, is indicated by the color purple.

To determine a putative list of TFs that may be involved in these reprogramming events, we performed a motif enrichment analysis (Figure 3G,H). The putative reprogramming TFs that act during PGC development include CTCF, Esrra, and Nrf1. Esrra is related to Esrrb, which, as mentioned, has been found to bind methylated DNA and de-methylate adjacent sequences. Although Nrf1 has been shown to be unable to bind methylated DNA, Nrf1 binding is often concurrent with the binding of other TFs nearby, such as CTCF, some of which can promote its interaction with DNA by promoting de-methylation of the Nrf1 binding site [36]. The fact that Nrf1 is the most highly enriched TF may reflect the need to be adjacent to a reprogramming TF binding site. Taken together, these results suggest that CpGs at binding sites for these TFs can become reprogrammed at specific stages of embryonic development. The TFs in Figures 3G and 3H therefore represent putative reprogramming TFs that are either capable of binding methylated DNA and promoting its de-methylation, or generally binding adjacent to a TF or other genomic feature that is reprogrammable.

### Reprogramming of CpG methylation status between ESCs and adult tissue occurs only at a small fraction of CpGs at binding sites for specific TFs

It has been shown that tissue-specific developmental and adult enhancers have highly methylated CpGs in epiblasts but low methylation levels in adult intestinal tissue, suggesting that DNA methylation at these sites is reprogrammed [37]. In order to determine the degree to which reprogramming between embryonic and adult somatic tissue occurs, we re-analyzed the data of Jadhav et al [37] in the context of individual distal CpGs. We chose to look at E7.5 embryos rather than E6.5 epiblast cells since they have globally higher methylation levels and similar methylation patterns, and thus make reprogramming easier to detect. About half of the individual CpGs at ATAC-seq peaks during intestinal development are highly methylated in E7.5 embryos but lowly methylated in the adult intestine (Additional file 1: Figure S3A). This is consistent with the original published results showing that CpGs at these tissue-specific enhancers are reprogrammed at some point after the epiblast stage [37]. In order to quantify the amount of global CpG methylation reprogramming that occurs during development to adulthood, we looked at the methylation levels in E7.5 embryos and adult intestine at E14.5m PGC DNase-Hi and DNase-Lo TF binding sites (Figure 4A; Additional file 1: Figure S3B). 70% of CpGs at E14.5m DNase-Lo sites with very high methylation levels (>80%) in E7.5 also have very high methylation levels in the adult intestine. On the other hand, 91% of E14.5m DNase-Hi sites with very low methylation (<20%) in E7.5 embryos also have very low methylation in the adult intestine. We performed the same analysis using BS-seq data from neonatal heart, kidney, and forebrain obtained from ENCODE, and obtained similar results (Additional file 1: Figure S3C-E). These results suggest that the methylation status of the majority of CpGs at TF binding sites remains consistent between late embryonic and adult tissue, but that some of them are reprogrammed. When we restricted the set of E14.5m PGC DNase-Hi and DNase-Lo sites to only those overlapping an ATAC-seq peak present after the epiblast stage, at either late embryonic, fetal, or adult stages, about half the CpGs at DNase-Lo sites with high E7.5 methylation have low methylation in the adult intestine (Figure 4B), recapitulating the results of Jadhav et al [37]. This is consistent with the hypothesis that CpGs bound by TFs after the epiblast stage tend to be highly methylated in the epiblast but then are reprogrammed to become functional during development or in adulthood, but that similar sequences are not reprogrammed when a TF does not bind during differentiation.

**Fig. 4.**
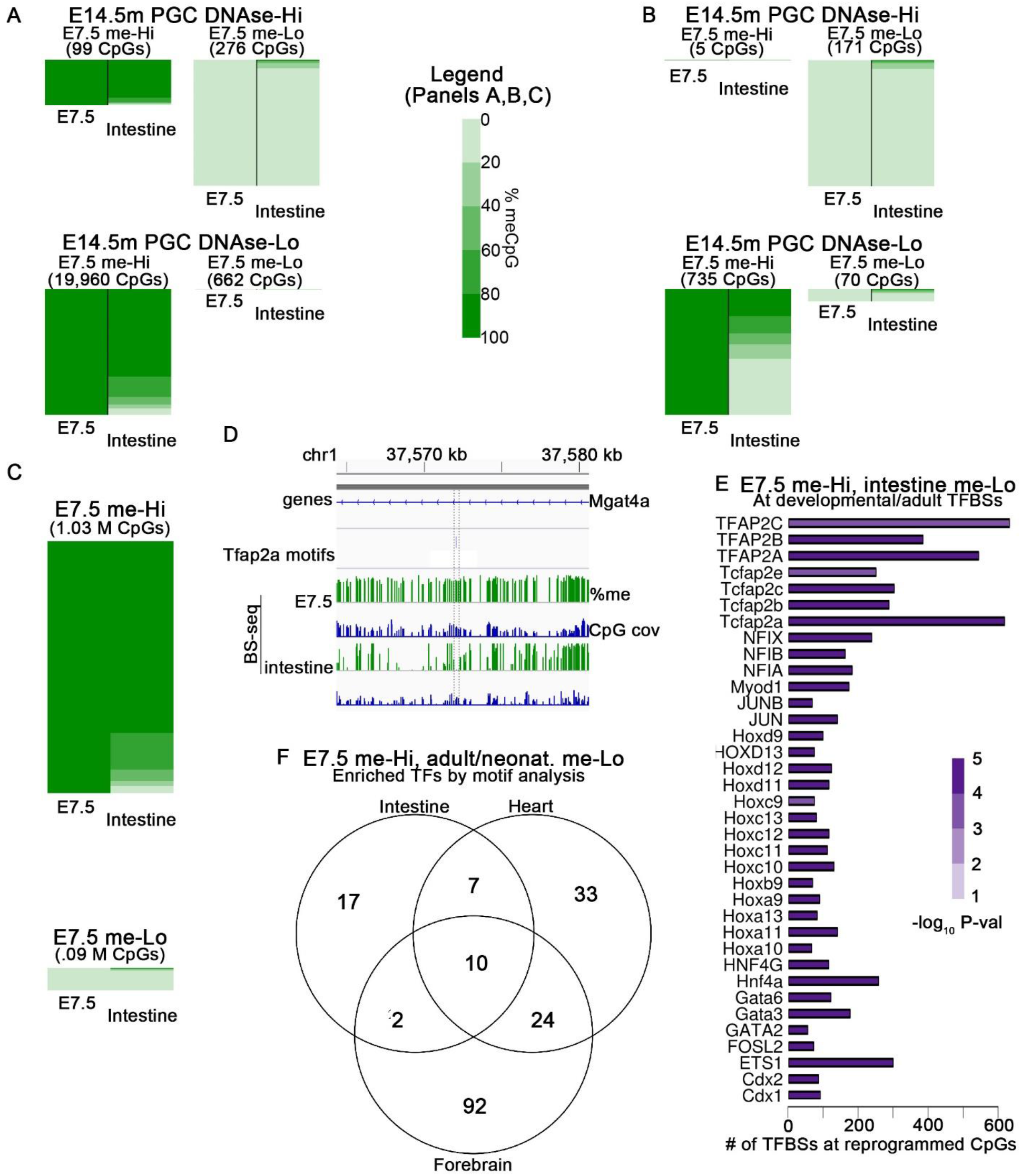
Reprogramming of CpG methylation status between ESCs and adult tissue occurs only at a small fraction of CpGs at binding sites for specific TFs. **A** Heatmap of CpG methylation percentage from BS-seq data in E7.5 embryos and adult intestine. Only CpGs with at least 10 BS-seq reads are displayed. The heatmaps are divided as follows: CpGs at E14.5m PGC DNase-Hi sites with very high methylation (> 80%) in E7.5 embryos, sorted by % methylation in intestine (top left); CpGs at E14.5m PGC DNase-Lo sites with very low methylation (< 20%) in E7.5, sorted by methylation in intestine (bottom left); CpGs at E14.5m PGC DNase-Hi sites with very high methylation in E7.5, sorted by methylation in intestine (top right); CpGs at E14.5m PGC DNase-Lo sites with very low methylation in E7.5, sorted by methylation in intestine (bottom right). The number of CpGs in each category is shown in the matrix to the right. The heatmaps at DNase-Hi sites are scaled separately from those at DNase-Lo sites to allow viewing of each class. The relative heights of the DNase-Hi heatmaps are scaled according to the relative number of CpGs at each, and the DNase-Lo hetmaps are scaled to each other similarly. **B** This heatmap is a subset of the sites in A, containing only the sites that overlap an ATAC-seq peak during embryonic, fetal, and adult intestinal development. **C** A heatmap of all CpGs genome-wide with at least 10 BS-seq reads on both samples that have either very high or very low methylation in E7.5 embryos. **D** An example of a region with a CpG at a binding sequence for the putative reprogramming TF Tcfap2a that has high methylation in E7.5 and low methylation in the adult intestine. **E** Bar plots similar to Fig. 3G,H, showing the number of motifs and the statistical significance of motif enrichment for the indicated TFs. **F** Venn diagram showing the overlap of sites that have high DNA methylation (>80%) in E7.5 embryos and low methylation (<20%) in either adult intestine, neonatal heart, or neonatal forebrain.

In order to determine if reprogramming at TF binding sequences reflects reprogramming of CpGs genome-wide, we examined all CpGs with very high and very low methylation in E7.5 and found that 76% of genome-wide CpGs with very high methylation in E7.5 also have very high methylation in the adult intestine, while 86% of CpGs with very low methylation in E7.5 also have very low methylation in adult intestine (Figure 4C). These are remarkably similar to the corresponding percentages when considering only TF binding sequences (Figure 4A), demonstrating that CpG methylation occurs only at a small percentage of CpGs genome-wide, or conversely that the methylation status of the majority of CpGs is maintained between embryos and adult somatic tissue. Figure 4D shows an example of a region around a TF biding sequence that is very highly methylated in E7.5 but that has lost its methylation in adult intestine. The fact that the percentage of reprogrammed CpGs increases drastically at TF binding sequences where a TF is known to bind at some point during the course of development and differentiation from embryos to the adult intestine suggests that this methylation reprogramming may be driven by the binding of reprogramming TFs to methylated DNA and promote their de-methylation.

A number of TFs widely studied in the context of cell differentiation, including Hox, Fos, and Jun family members, are significantly enriched at sites that are reprogrammed between E7.5 embryos and the adult intestine and that bind a TF at some point during the course of development and differentiation into adult intestinal cells (Figure 4E). We hypothesize that at least some of these TFs are capable of binding to methylated DNA and subsequently recruiting Tet enzymes to de-methylate nearby CpGs. A similar analysis of neonatal heart, kidney, and forebrain samples from ENCODE reveals sets of TFs with both overlapping and distinct TFs to those of intestine (Figure 4F; Additional file 1: Figure S3F-H). This suggests that, over the course of development, CpGs that are methylated in the embryo become reprogrammed in a tissue-specific manner. Our results suggest that specific reprogramming TFs are likely to be expressed only at specific stages and within specific cell lineages, forming the basis for cell-specific enhancers and other regulatory elements. However, our results suggest that the methylation status of the majority of CpGs in the genome is preserved from embryonic to adult tissues.

### TF binding site affinity influences the binding patterns of TFs during PGC and embryonic development

We note that in our analyses up to this point, for each of the TFs expressed in PGCs and ESCs, only a fraction of their putative binding sites in the genome are actually bound by a TF. Except for Intracisternal A Particles (IAPs), CpGs are largely unmethylated in E13.5m PGCs [3]. In the absence of DNA methylation as an epigenetic determinant (or at least an indicator of an epigenetic determinant) of TF binding in E14.5m PGCs and ESCs, we hypothesize that TFs would bind more frequently to high affinity DNA sequences than to low affinity ones. To test this, we employed Transcription Factor Affinity Prediction (TRAP), a tool that quantifies TF affinity for a given sequence using biophysical models [38]. TRAP has been shown to accurately predict the most likely TF to bind at a given set of regions [39]. We determined a TRAP affinity score for all peaks overlapping DNase-Hi and DNase-Lo sites in both E14.5m PGCs and ESCs, separately. In both cases, the median TRAP affinity was significantly higher in the set of DNase-Hi sites than the DNase-Lo sites (Figure 5A). The difference between DNase-Hi and DNase-Lo sites in ESCs was not as large as in PGCs. It is possible that this is due to the fact that global CpG methylation levels are lower in E13.5m PGCs than in ESCs and therefore more CpGs contain epigenetic information in ESCs than in E13.5m PGCs that can be regulated by factors besides DNA sequence binding affinity. We next combined all TF binding sequences as determined by fimo, not just DNase-Hi and DNase-Lo sites, and divided them according to their TRAP affinity score. The average DNase-seq signal was significantly higher in regions with high TRAP affinity than regions with low TRAP affinity in both E14.5m PGCs and ESCs (Figure 5B).

**Fig. 5.**
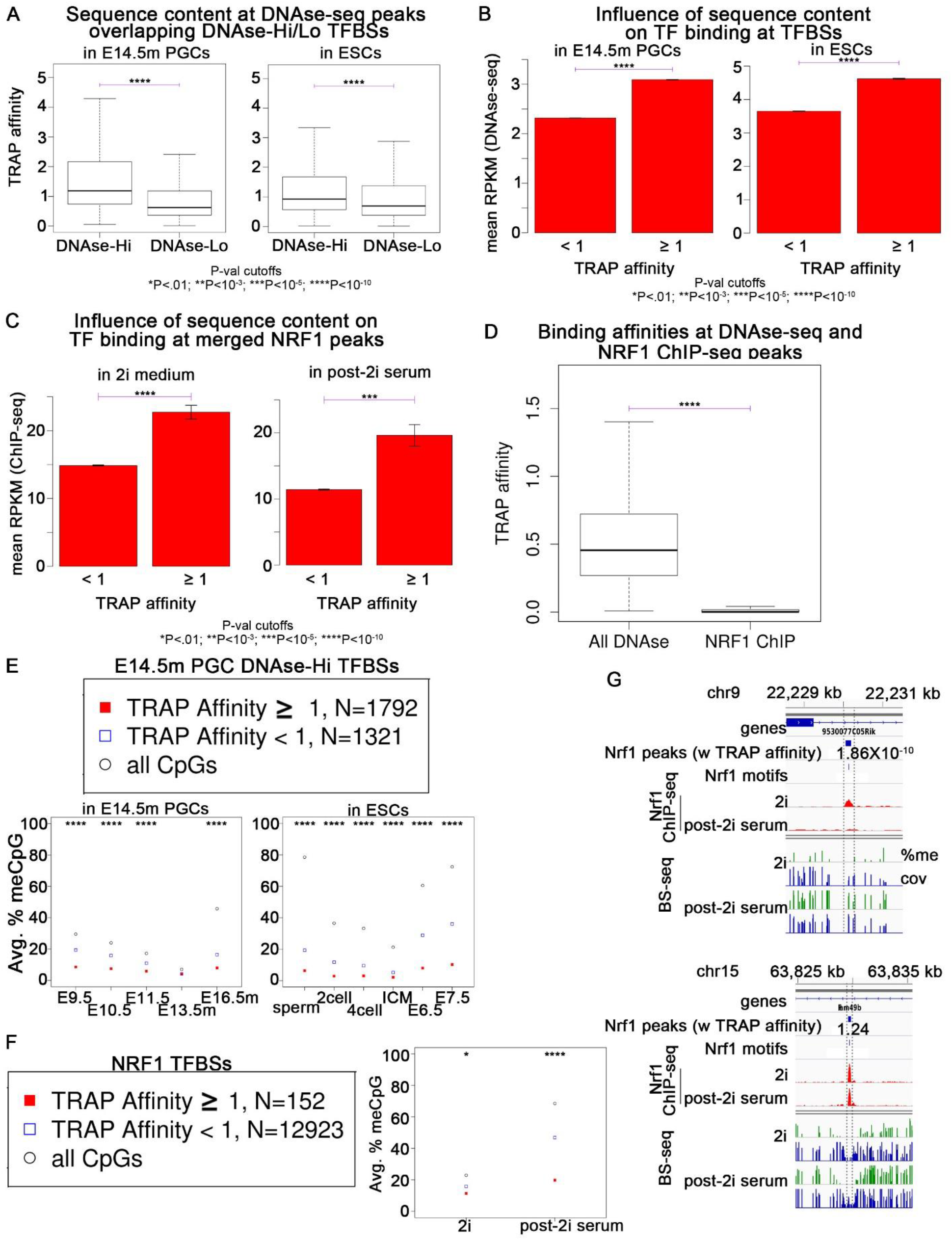
TF binding site affinity influences the binding patterns of TFs during PGC and embryonic development. **A** Boxplots showing the distribution of TRAP affinity scores for embryonic TFs at E14.5m PGC DNase-Hi and DNase-Lo sites. P-values are by the Wilcoxon rank-sum test. **B,C** Average DNase-seq signal at regions with high and low TRAP affinities in the indicated samples. P-values are by the student’s t-test. **D** Boxplot comparing TRAP affinity at Nrf1 ChIP-seq peaks to those of DNase-seq peaks. P-values are by the Wilcoxon rank-sum test. **E,F** Weighted average DNA methylation levels in the indicated samples at sites with high versus low TRAP affinity scores for PGC TFs at E14.5m PGC DNase-seq peaks (E) and for Nrf1 at Nrf1 ChIP-seq peaks (F). **G** Example showing the Nrf1 ChiP-seq and DNA methylation levels during and after growth in 2i medium at a Nrf1 peak with a relatively high TRAP affinity.

It has been shown that binding of Nrf1 in ESCs grown in 2i medium, which results in global CpG demethylation, can be outcompeted by DNA methyltransferases and become re-methylated after removal from 2i medium [36]. These results could be explained if Nrf1 has a relatively low binding affinity for DNA compared to other TFs. To test this, we computed TRAP affinity scores at all Nrf1 peaks during and after 2i medium exposure. The average ChIP-seq signal was significantly higher at high-affinity peaks in both 2i and standard media (post-2i) (Figure 5C), consistent with previous TRAP affinity results [39]. Indeed, the median TRAP affinity at NRF1 peaks was significantly lower than the median TRAP affinity of E14.5m PGC DNase-seq peaks (Figure 5D). To further test the hypothesis that high affinity TF binding sites inhibit de-novo DNA methylation better than low affinity sites, we divided the E14.5m PGC and ESC DNase-Hi sites into those with high and low TRAP affinity, and found that DNA methylation levels were significantly lower at sites with high affinity than at those with low affinity in both PGCs and ESCs (Figure 5E). Similarly, we divided Nrf1 ChIP-seq peaks into those with high and low TRAP affinities for Nrf1, and found that after removal from 2i medium, high-affinity Nrf1 binding sites had significantly lower DNA methylation than low-affinity Nrf1 sites (Figure 5F,G). Taken together, these results suggest that high affinity binding sites, as determined by TRAP, for TFs expressed in PGC and ESC development will be bound more frequently by their TFs than low affinity sites, and that sites with high affinity will be protected from de-novo DNA methylation to a greater extent than low affinity sites in both PGCs and ESCs.

### IAPs possess a relatively low affinity for embryonic reprogramming TFs and high affinity for non-reprogramming TFs

IAPs are the main class of DNA sequences that remain methylated in E13.5m PGCs [3]. Based on the last section, we hypothesized that the DNA binding domain of reprogramming TFs present in PGCs would have a relatively low binding affinity for typical IAP sequences. To test this, we performed a motif enrichment for TFs within annotated IAP Long Terminal Repeats (LTRs) that are expressed in E16.5m PGCs (Figure 6A). Some TFs associated with reprogramming (Figures 3G,H) are present in the list of significantly enriched TFs at IAP LTRs (Figure 6A). We therefore sought to compare the affinity of TFs associated with reprogramming to TFs not associated with reprogramming, limiting the analysis to only those TFs expressed during global DNA re-methylation in E16.5m PGCs. For each IAP LTR, we determined putative reprogramming TFs expressed in E16.5m that have the highest TRAP affinity, and used this information as the reported reprogramming affinity for that TF binding site. We similarly determined which non-reprogramming TFs expressed in E16.5m have the highest affinity and used its affinity as the reported affinity for that TF binding site. We then compared the distribution of these values over all annotated IAP LTRs and found that the median affinity of putative reprogramming-associated TFs was significantly lower than the median affinity for non-reprogramming TFs (Figure 6B), supporting the hypothesis that IAPs have an inherently low affinity for TFs that can bind to methylated DNA and assist in their de-methylation.

**Fig. 6.**
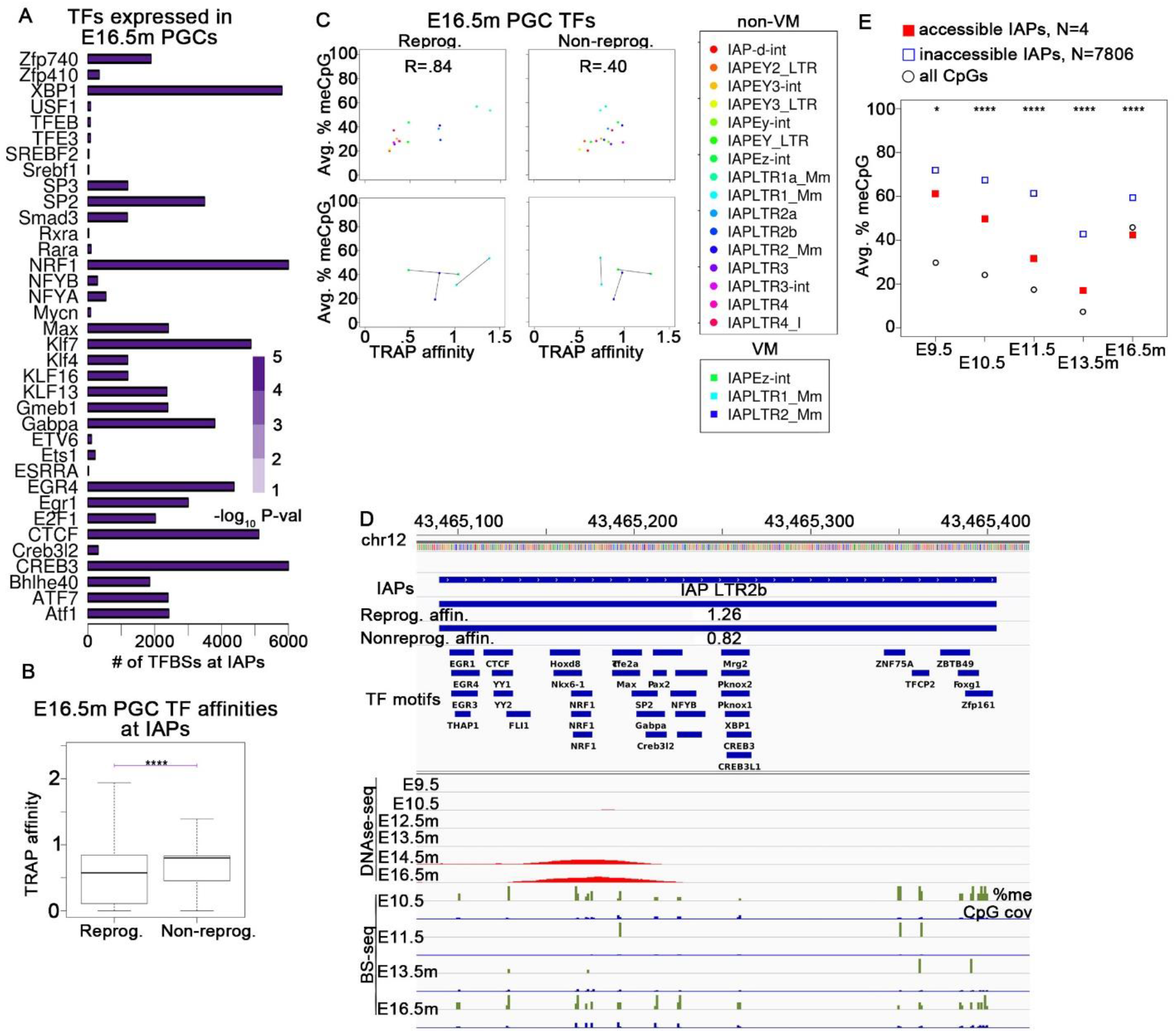
IAPs possess a relatively low affinity for PGC reprogramming TFs and high affinity for non-reprogramming PGC TFs. **A** Bar plots showing the motif enrichment of TFs expressed in E16.5m PGCs at IAP LTR regions. **B** Box plots comparing the TRAP affinity of TFs with evidence of DNA methylation reprogramming in PGCs versus TFs expressed in PGCs with no evidence of DNA methylation reprogramming capabilities. **C** Scatterplots comparing weighted-average CpG methylation in E13.5m PGCs (y-axis) to TRAP affinities (x-axis) of putative reprogramming TFs expressed in E16.5m PGCs (left column) as well as non-reprogramming TFs expressed in E16.5m PGCs (right column). The top row of scatterplots consists only of non-VM IAPs while the bottom row of scatterplots compares VM-IAPs to non-VM-IAPs, where a line is drawn between values from VM-and non-VM-IAPs of the same class. **D** Example showing DNase-seq and DNA methylation levels during PGC development at an IAP LTR that has evidence of TF binding. Tracks showing the highest TRAP affinity of the E16.5m PGC putative reprogramming TFs, and of E16.5m PGC non-reprogramming TFs, at the whole IAP, are displayed under the IAP track. **E** Average DNA methylation levels at DNase-seq accessible vs. inaccessible IAPs. The following P-value cutoffs apply to B and E: *P<.01; **P<.001; ***P<10^−5^;****P<10^−10^.

Our results suggest that low affinity for reprogramming TFs may be a contributing factor in helping to ensure that IAPs resist global de-methylation. To further characterize the degree of contribution of DNA sequence of IAPs to their demethylation resistance, we plotted the average CpG methylation at E13.5m PGCs versus the average binding affinity for reprogramming and non-reprogramming TFs expressed in E16.5m PGCs for each class of annotated IAPs in mouse (Figure 6C). Surprisingly, there is a strong positive correlation (0.84) between CpG methylation levels in E13.5m PGCs and the affinity for putative reprogramming TFs, whereas the correlation of E13.5m PGC CpG methylation with non-reprogramming TF affinities is much weaker (0.40). This shows that the greater the affinity an IAP has for TFs associated with CpG methylation reprogramming, the greater its resistance to de-methylation during PGC development. This observation suggests that the mechanism of IAP de-methylation resistance is sensitive to the affinity of TFs associated with DNA methylation reprogramming and that one function of this mechanism may be to protect IAPs from binding to reprogramming TFs.

Certain IAPs with variable DNA methylation across individuals but persistent methylation across tissues of an individual are known as Variably Methylated IAPs (VM-IAPs) [1]. In order to assess the role of TF binding and de-methylation resistance in the formation of VM-IAPs, we compared the average CpG methylation in E13.5m PGCs and the affinity of putative reprogramming and non-reprogramming TFs expressed in E16.5m PGCs at VM vs non-VM IAPs of the same class, for the 3 classes of IAPs with at least 25 validated VM-IAPs (Figure 6C). Interestingly, the TRAP affinity of reprogramming TFs at VM-IAPs appears close to 1 for all 3 classes, and classes of IAPs that have an affinity close to 1 at non-VM IAPs change very little between their VM counterparts, whereas IAPs whose non-VM affinity is smaller than 1 increase to near 1 in their VM counterparts. Furthermore, IAPs whose non-VM IAPs have a reprogramming TF affinity larger than 1 decrease to near 1 in their VM-counterparts. Such a relationship does not exist for non-reprogramming TFs (Figure 6C). These results suggest that VM-IAPs are more likely to form when they have an intermediate affinity for a reprogramming TF. This may be due to observations shown in Figure 6B: since IAPs with a very high affinity for reprogramming TFs are highly resistant to de-methylation in PGCs, their high level of de-methylation resistance may be sufficient to escape methylation reprogramming, whereas IAPs with moderate affinity for reprogramming TFs will have moderate DNA-methylation resistance in PGCs. Therefore, there may be a stochastic competition between reprogramming and de-methylation resistance in PGCs that results in VM at intermediate values of each. Experimental analyses will be required to test this hypothesis. While no obvious relationship between the emergence of VM and the affinity for non-reprogramming TFs emerged in our analysis, their affinities did change between VM and non-VM IAPs of the same type, suggesting that the final methylation state at IAPs in adults may arise out of a complex interplay between the affinity of reprogramming and non-reprogramming TFs and de-methylation resistance in PGCs and ESCs.

We next looked for IAPs accessible to DNase-seq in PGCs. Although rare, four annotated IAPs were accessible to DNase-seq, and these have a relatively high affinity for putative reprogramming TFs (Figure 6D; Additional file 1: Figure S4A,B). There are likely more than just four TF-bound IAPs genome-wide, but the mappability at IAPs with this DNase-seq data was relatively low since they were obtained by single-end sequencing, while the paired-end BS-seq data is clearly more mappable (Figure 6D; Additional file 1: Figure S4B,C). In any event, this demonstrates that TFs expressed during PGC development are capable of binding to IAP LTR sequences. Our results suggest that this would result in a trans-generational escape from DNA hypermethylation at the IAP if one or more TFs constitutively present in the nucleus can bind to high-affinity sites during PGC and ESC development after some event interferes with de-methylation resistance. Indeed, the IAPs accessible to DNase-seq in PGCs have a significantly and substantially lower DNA methylation level throughout PGC development across the entire LTR compared to IAP LTRs that are not accessible to DNase-seq in E14.5m PGCs (Figure 6E). These results suggest that if the affinity of an IAP for a PGC/ESC reprogramming TF can be increased, either by sequence alterations or perhaps by environmentally-induced over-expression of a reprogramming TF, it can escape persistently high DNA methylation levels across generations, resulting in a trans-generational change in epiphenotypes.

## Discussion

Key observations in this study include the fact that the DNA methylation status is faithfully preserved after global de-methylation and re-methylation at the majority of CpGs in the mouse genome, and that whether a CpG is methylated or not before and after global de-/re-methylation is highly correlated with TF binding in PGCs and ESCs during re-methylation. Although it is commonly suggested that trans-generational epigenetic inheritance can only occur through CpGs that resist de-methylation during PGC and ESC differentiation, results described here suggest that this process may occur even in the absence of de-methylation resistance, because TF binding during the re-methylation phases of PGC and ESC development preserve the epigenetic information at their binding sites. These results support models of trans-generational epigenetic inheritance that do not require DNA de-methylation resistance, effectively increasing the number of CpGs through which these phenomena can occur from 10% to an estimated 78%. At non-IAP sites or other CpGs that resist global DNA de-methylation during PGC and preimplantation development, CpGs present in high-affinity TF binding sites that are unmethylated during germ line development will be able to bind their cognate TFs and will subsequently be protected from de-novo methylation (Figure 7A). Conversely, non-IAP sites with low affinity for TFs will not be able to bind TFs during PGC and ESC differentiation and will not be protected from de-novo methylation. The methylation state of such sites will be persistently high between generations, except during the global de-methylation phases of PGC and pre-implantation development, after which it will be restored and maintained trans-generationally (Figure 7A).

**Fig. 7.**
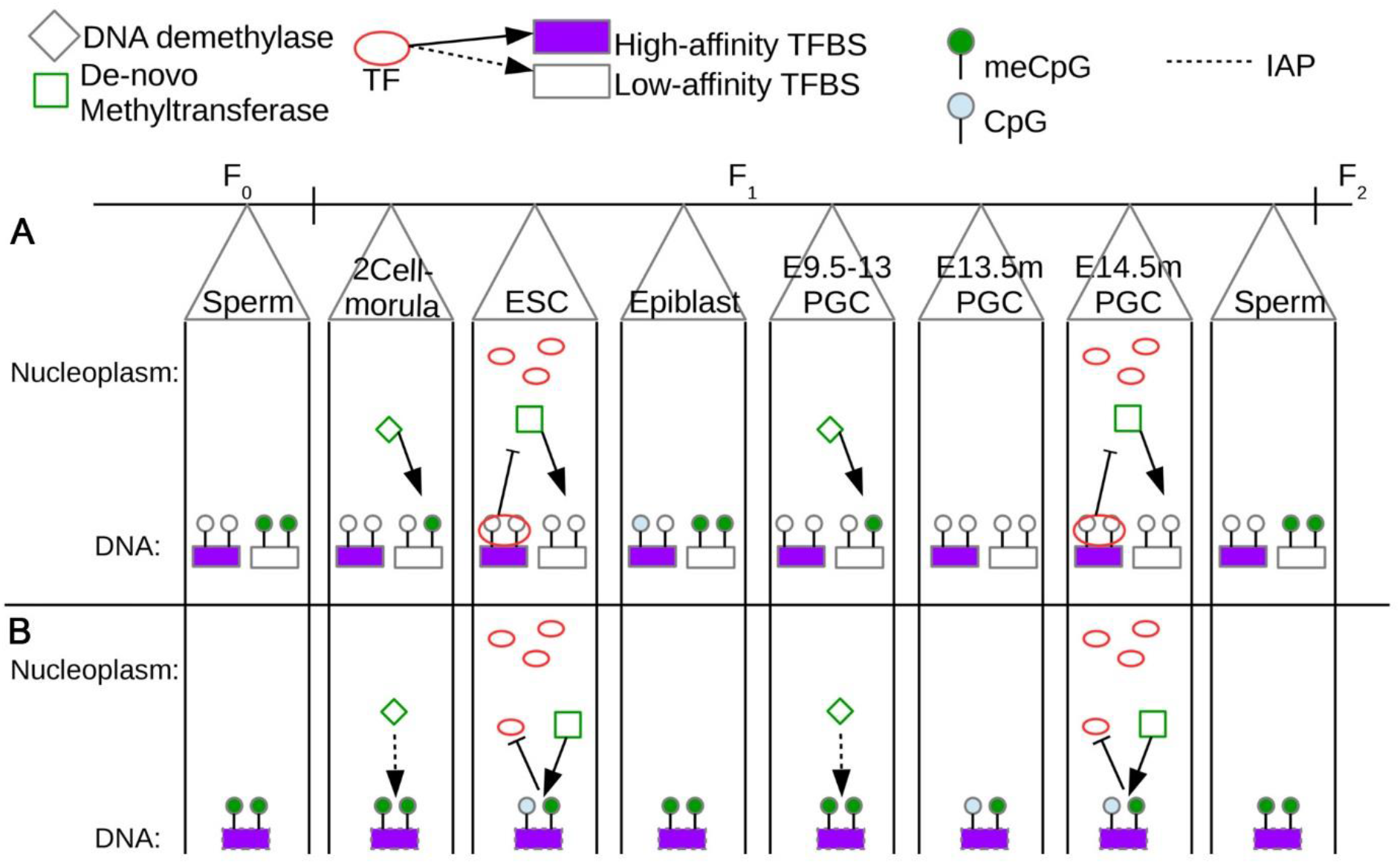
Transcription factor mediation of trans-generational epigenetic inheritance in mammals. **A** This panel follows the methylation status of two CpGs at 2 different genomic loci which are not resistant to global de-methylation in PGCs: one is a TF binding site with a high affinity for a TF present in E14.5m PGCs as well as ESCs, while the other locus has a low affinity for such TFs. The high affinity site is perpetually hypomethylated because the TF binds during the global re-methylation phases of PGC and ESC development, whereas the low affinity TF binding site is perpetually hypermethylated at all stages except for the globally hypomethylated stages of PGC and ESC differentiation. Because a TF cannot bind with high affinity during global re-methylation, the low affinity TF binding site will become re-methylated and will be hypermethylated at all other developmental stages in the absence of binding of a reprogramming TF. **B** This panel shows the behavior for a pair of CpGs at a site with high affinity for a PGC and ESC TF that is within an IAP or any other genomic locus that is resistant to de-methylation during PGC and pre-implantation development. Because the CpG escapes global de-methylation, it never binds the TF and remains perpetually hypermethylated. In E14.5m PGCs and ESCs, the methylation levels are reduced, but never become completely unmethylated, such that during the global re-methylation phase, they become hypermethylated once again. CpGs at low affinity TF binding sites within de-methylation resistant loci will behave in the same manner as high affinity TF binding sites, except that the high affinity sites will have a higher likelihood of binding their TF if the de-methylation resistant machinery can be interfered with in some way.

During adult somatic differentiation, regions that are methylated in the epiblast may become de-methylated at binding sites for specific TFs in order to define developmental and tissue-specific enhancer regions. However, the majority of CpGs genome-wide follow the usual cycle of de-methylation and re-methylation, and remain methylated until the next generation. At IAPs or other sites that resist global PGC/pre-implantation de-methylation, CpGs remain persistently hypermethylated throughout development across generations whether or not they have high affinity for non-reprogramming TFs expressed during the remethylation phases of PGC and embryonic differentiation (Figure 7B). IAPs are the only class of DNA sequences shown to escape de-methylation followed by re-methylation during PGC development in mice, although de-methylation resistance has been also observed at a small number of non-IAP CpGs [3]. Consistent with persistent hypermethylation at IAPs, TFs rarely bind to IAPs - even at high-affinity binding sites - in the germ line (Figure 7B).

An environmental stimulus capable of altering the ability of IAPs or other DNA sequences to resist de-methylation during PGC and pre-implantation development would cause a switch to the normal pattern of either persistent hypomethylation, or de-methylation followed by re-methylation, depending on the binding of TFs in PGCs and ESCs. This is equivalent to changing from behaving as in Figure 7B to behaving as in Figure 7A. This could have trans-generational effects if the loss of de-methylation resistance persists across generations. An obvious way in which this could happen is through a mutation that ablates the de-methylation resistance but still leaves a high affinity for a TF to bind prior to global re-methylation.

Alternatively, if the de-methylation resistance could be blocked epigenetically, a transgenerational effect would be observed as long as the epigenetic inhibition of the ability to block de-methylation persists. Our results suggest that if a TF binds to an IAP it will interfere with de-methylation resistance, so any epigenetic event that can allow a TF to bind to a region capable of resisting global de-methylation would result in a change in molecular epiphenotypes. We found that IAPs have a low affinity for TFs associated with DNA methylation reprogramming during PGC development, which suggests that overexpressing such TFs at the appropriate stage of PGC development may allow it to bind long enough to de-methylate CpGs in its vicinity on the IAP. At this point, other non-reprogramming TFs could bind and then help the IAP escape persistent hypermethylation, possibly trans-generationally, depending on the developmental expression patterns of the specific TFs that can bind nearby.

While the mechanisms of DNA de-methylation resistance are not fully understood, it is thought to involve piRNAs at IAPs [40]. IAPs are partially de-methylated during PGC and preimplantation development, suggesting that the mechanism of IAP DNA methylation maintenance is in competition with global de-methylation rather than completely ablating it. miRNAs and tRNAs both have the potential to regulate retroelements such as IAPs by affecting the translation of proteins involved in their regulation, and alteration of these RNAs has been linked to heritable epigenetic changes [10, 11, 13]. This suggests that one possible mechanism by which ncRNAs can induce inter- or trans-generational epigenetic effects, as has been observed, is by interfering with the ability of IAPs to resist de-methylation during PGC and preimplantation development. An increase in miRNAs could inhibit the translation of proteins needed to maintain methylation at IAPs, while a loss of tRNAs could have the same effect. Indeed, alterations in piRNAs have been observed in studies of transgenerational inheritance of stress-induced epiphenotypes [10], which as discussed, are thought to regulate DNA methylation at IAPs and other transposable elements. Thus, the alterations in miRNA’s are coincident with likely alterations in de-methylation resistance. In addition, sperm tRNAs are involved in downregulating expression of the MERVL transposable element in pre-implantation embryos in studies in studies showing the involvement of tRNA fragments in a low protein diet-induced trans-generational epiphenotypes [13]. These studies also observed that the level of several piRNAs were altered in sperm but did not explore this observation further. Nonetheless, these studies establish a link between ncRNAs and regulation of transposable element de-methylation resistance in inter- and trans-generational epigenetic inheritance [13].

There is evidence that specific histone variants such as H2A.Z may also act as mediators that maintain the DNA methylation status of CpGs across embryonic development [17]. Positioned nucleosomes containing specific histone variants could be an independent mechanism of maintaining the memory of previous DNA methylation states, or, alternatively, TFs and placeholder nucleosomes could act in concert to maintain epigenetic information during global de-methylation and re-methylation, as it has been shown that binding of pioneer TFs can direct nucleosome positioning [41]. Although we focused specifically at the actual TF binding sites in our analyses, all of the examples in this study show that the associations of TF binding and CpG methylation patterns across development still exists not just at the core binding site, but at the surrounding region as well. This is consistent with previous studies showing that knocking out two specific TF binding sites within a super enhancer results in increased de-novo DNA methylation of the entire super enhancer region [42]. Taken together, the evidence suggests that TF binding and placeholder nucleosome occupancy may be coupled, but further studies are needed to verify the existence of placeholder nucleosomes in mammals and the relationship of their localization to TF binding during PGC and pre-implantation development.

## Conclusion

The results of this study support models explaining the occurrence of inter- and trans-generational epigenetic inheritance in the absence of DNA de-methylation via TF-binding during global DNA re-methylation. Our results explain the preservation of CpG methylation status despite global de-methylation and re-methylation during PGC and pre-implantation development. These models predict that if de-methylation resistant genomic loci gain the ability to bind TFs in PGCs but not ESCs, or vice-versa, this would only result in inter- rather than trans-generational inheritance because it would only be protected from re-methylation during one of two rounds. In the round where sequences are not protected from re-methylation, these sites would become re-methylated and revert back to its original methylation status. Our model therefore provides a framework for the design of experiments intended to elucidate specific mechanisms of inter- and trans-generational epigenetic inheritance.

## Methods

### DNase-seq and ChIP-seq analysis

Raw fastq files were downloaded from the sources cited in the text and trimmed using Trimmomatic-0.38 [43] with the parameters “1:0:2 TRAILING:20 MINLEN:20”. Trimmed reads were mapped with bowtie [44] with the parameters -m 1 --mapq 254. Duplicate reads were then removed using MarkDuplicates from picard-tools version 2.1.0 (https://broadinstitute.github.io/picard/). For displaying in IGV, macs2 [16] predictd was used to determine the average fragment length, and mapped reads were shifted by half the average predicted fragment length towards the fragment center. The genome-wide, normalized coverage was then determined using bedtools [45] genomecov on the deduplicated, shifted reads, scaled by the total number of processed reads per million in each sample.

### ATAC-seq analysis

Raw fastq files were downloaded from the sources cited in the text and trimmed using pyadapter_trim.py (https://github.com/kundajelab/training_camp/blob/master/src/pyadapter_trim.py). To adjust the fragment size for transposase insertions, we aligned all reads as + strands offset by +4 bp and -strands offset by 5 bp [46]. Trimmed and offset reads were aligned to the mm9 reference genome using bowtie2 [47] with the parameter -X 2000. Only paired-end reads with fragment length between 50 and 115 bp, corresponding to TF binding sites [46], referred to in this paper as TF-ATAC, were kept. For viewing in IGV, a bed file of processed TF fragments was created, and the genome-wide, normalized coverage was then determined using bedtools genomecov, scaled by the total number of processed TF reads per million in each sample.

### BS-seq analysis

For the embryonic data [4], processed files containing read count information of methylated and unmethylated CpGs were downloaded and converted from mm10 to mm9 using liftover [48]. For all other BS-seq data, raw fastq files were downloaded and trimmed as described for the DNase-seq and ChIP-seq data. Trimmed reads were then aligned to the mouse mm9 reference genome using bismark [49] v0.19.0, deduplicated with deduplicate_bismark, and then CpG methylation was extracted using bismark_methylation_extractor. For each CpG, the number of meCpG and CpG reads for all replicates of a given sample were combined to obtain a single average methylation value per sample per CpG. For heatmaps, the R function pheatmap was used, and only CpGs with > 10 BS-seq reads were used. For plots of average methylation, the average was calculated using all CpGs, with each CpG being weighted by the total number of BS-seq reads at that CpG. This is equivalent to simply pooling all BS-seq reads at the CpGs being considered at taking the overall average.

### RNA-seq analysis

Raw fastq files were downloaded from the sources cited in the text, and reads were trimmed as described in the “DNase-seq and ChIP-seq analysis” section. Reads were then aligned to the mm9 reference genome using tophat2 [50] with the parameters --no-mixed --no-discordant, and non-uniquley mapped reads were discarded. FPKM values for annotated genes were calculated using cuffdiff.

### Estimation of the percentage of CpGs with a preserved methylation status prior to and after global de-methylation and re-methylation of PGCs and preimplantation embryos

Only 9% of CpGs in sperm, or 0.15 million CpGs, had intermediate methylation levels (between 20 and 80%), and we conservatively assume that none of those will be preserved before and after PGC and preimplantation reprogramming. Of the 1.38 million CpGs with > 80% methylation in sperm, 91% had > 50% methylation in epiblasts, but since the epiblast is still undergoing de-novo methylation, we assume that anything over 50% in epiblast should eventually become over 80% methylated in sperm and so we consider that these are CpGs whose methylation status is preserved before and after PGC and preimplantation reprogramming. The remaining 0.18 million CpGs in sperm have < 20% methylation, and of these, 46% also have < 20% methylation in epiblasts, so we consider that 46% are preserved. Based on these assumptions, the percentage of preserved CpGs is given by the weighted average of the percentages of preserved CpGs out of those that are very high in sperm, intermediate in sperm, and very low in sperm, as described above. Explicitly, the calculation is as follows: (1.38×0.91+0.18×46 + 0.15×0)/(1.38+0.18+0.15).

### Estimation of TF binding affinities based on DNA sequence

Transcription Factor Affinity Prediction [38] scores were calculated using TEPIC [51] at PGC DNase-seq peaks downloaded from NCBI GEO [22] as well as at annotated IAP LTRs. An affinity score was calculated for all TFs in the “Merged_JASPAR_HOCOMOCO_KELLIS_Mus_musculus.PSEM” file provided with the TEPIC software. For each region where TRAP affinities were calculated, the maximum TRAP affinity of the TFs under consideration was chosen as the affinity score used for that region in subsequent analyses, since in PGCs, at any given TFBS for an embryonic TF, the TF with the highest binding affinity is likely to be the one that binds to that region.

## Supplementary information

Supplementary information accompanies this paper.

## Additional file 1

**Figure S1.** Related to Figure 1. **Figure S2.** Related to Figure 2. **Figure S3.** Related to Figure 4. **Figure S4.** Related to Figure 6.

## Additional file 2

Table S1.

## Authors’ contributions

IK and VGC conceived and designed the study. IK performed all the analyses. IK and VGC wrote and approved the manuscript.

## Availability of data and materials

All datasets analyzed in the present study are available in either the NCBI GEO, ENCODE, or European Nucleotide Archive repositories at the following locations. Epiblast and PGC DNase-seq GSE109770; PGC BS-seq, PRJEB3376: Preimplantation DNase-seq, GSE76642; Preimplantation and Sperm BS-seq, GSE56697; RS ATAC-seq, GSE102954; Sperm ATAC-seq, GSE116857; ICM ATAC-seq, GSE66390; ESC ATAC-seq, GSE67299; Adult intestine ATAC-seq and BS-seq, GSE111024; Neonatal forebrain BS-seq, GSE82356; Neonatal heart BS-seq, GSE82658; Neonatal kidney BS-seq, GSE82451; NRF1 ChIP-seq and BS-seq in 2i and standard serum, GSE67867; Mouse DNase-seq peaks, http://hgdownload.soe.ucsc.edu/goldenPath/mm9/encodeDCC/wgEncodePsuDnase/; Mouse DNase-seq peaks, http://hgdownload.soe.ucsc.edu/goldenPath/mm9/encodeDCC/wgEncodeUwDnase/;.

Processed data or scripts to generate processed data used in this report are available upon request.

## Ethics approval and consent to participate

Not applicable

