## Supplemental Figures for "Transcription factors protect from DNA re-methylation during reprograming of primordial germ cells and pre-implantation embryos"

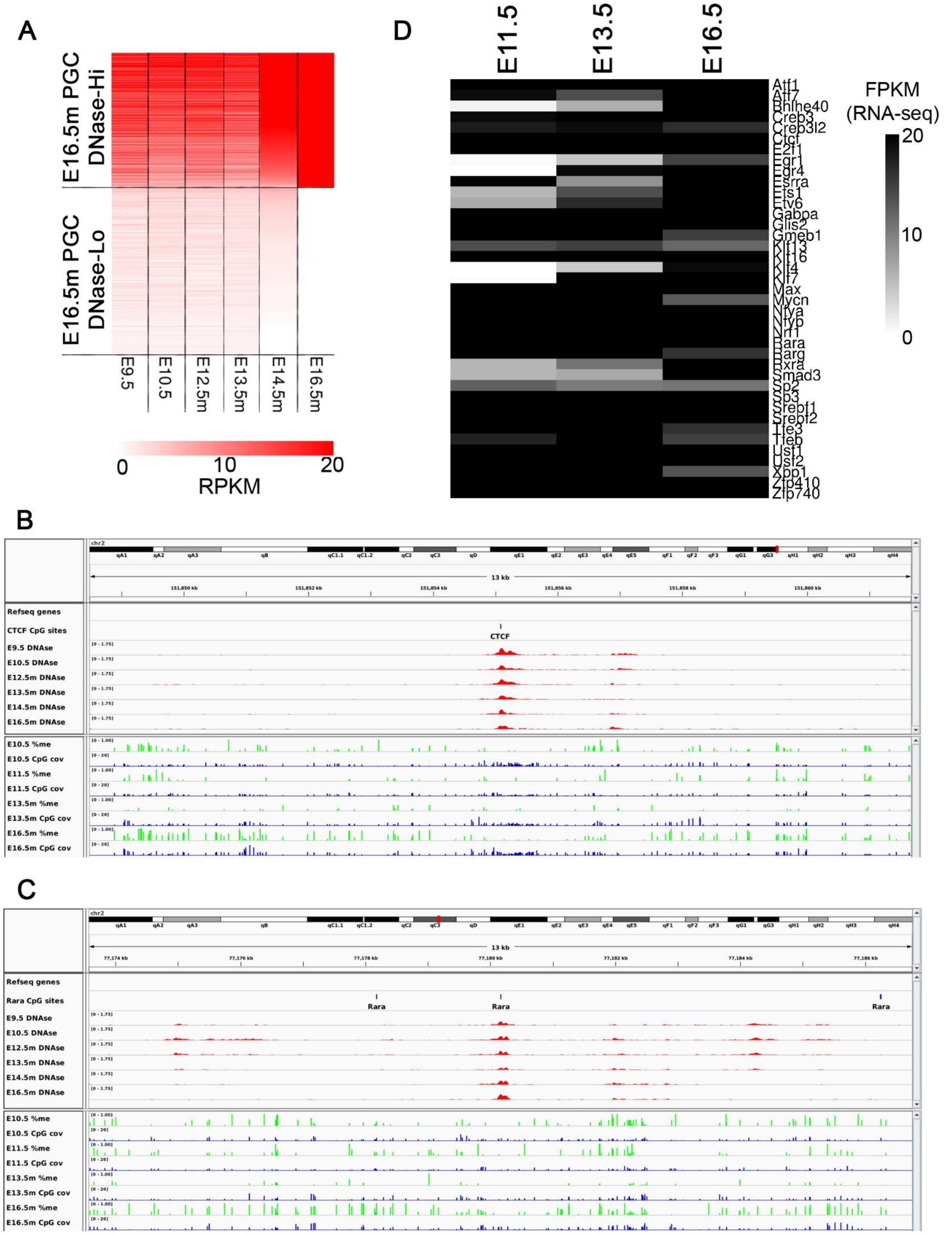
A
E16.5m PGC DNase-Hi
E16.5m PGC DNase-Lo
E9.5
E10.5
E12.5m
E13.5m
E14.5m
E16.5m
0
10
20
RPKM
D
E11.5
E13.5
E16.5
Atf1
Atf7
Bhlhe40
Creb3
Creb3l2
Ctcf
E2f1
Egr1
Egr4
Esrra
Ets1
Etv6
Gabpa
Glis2
Gmeb1
Klf13
Klf16
Klf4
Klf7
Max
Mycn
Nfya
Nfyb
Nrf1
Rara
Rarg
Rxra
Smad3
Sp2
Sp3
Srebf1
Srebf2
Tfe3
Tfeb
Usf1
Usf2
Xbp1
Zfp410
Zfp740
FPKM
(RNA-seq)
20
10
0
B
chr2
13 kb
Refseq genes
CTCF CpG sites
CTCF
E9.5 DNAse
E10.5 DNAse
E12.5m DNAse
E13.5m DNAse
E14.5m DNAse
E16.5m DNAse
E10.5 %me
E10.5 CpG cov
E11.5 %me
E11.5 CpG cov
E13.5m %me
E13.5m CpG cov
E16.5m %me
E16.5m CpG cov
C
chr2
13 kb
Refseq genes
Rara CpG sites
Rara
Rara
Rara
E9.5 DNAse
E10.5 DNAse
E12.5m DNAse
E13.5m DNAse
E14.5m DNAse
E16.5m DNAse
E10.5 %me
E10.5 CpG cov
E11.5 %me
E11.5 CpG cov
E13.5m %me
E13.5m CpG cov
E16.5m %me
E16.5m CpG cov

**Figure S1.** Related to Figure 1. **A** Heatmap of RPKM values within just binding sequences called by fimo, separated into those that have high DNase signal in E16.5m PGCs (DNase-Hi, RPKM > 20) and those that have RPKM=0 (DNase-Lo). Rows are ordered by decreasing signal in E14.5m. All stages shown are during PGC development. **B,C** Example region with DNase accessibility and low local DNA methylation throughout PGC development. CpG cov indicates the BS-seq read coverage. TF binding sequences called by fimo that overlap the peak summit in E14.5m are shown. **D** RNA-seq levels in the indicated stages of PGC development for TFs that are expressed in E16.5m PGCs and were significantly enriched in a motif analysis at E14.5m PGC DNase-Hi sites.

A

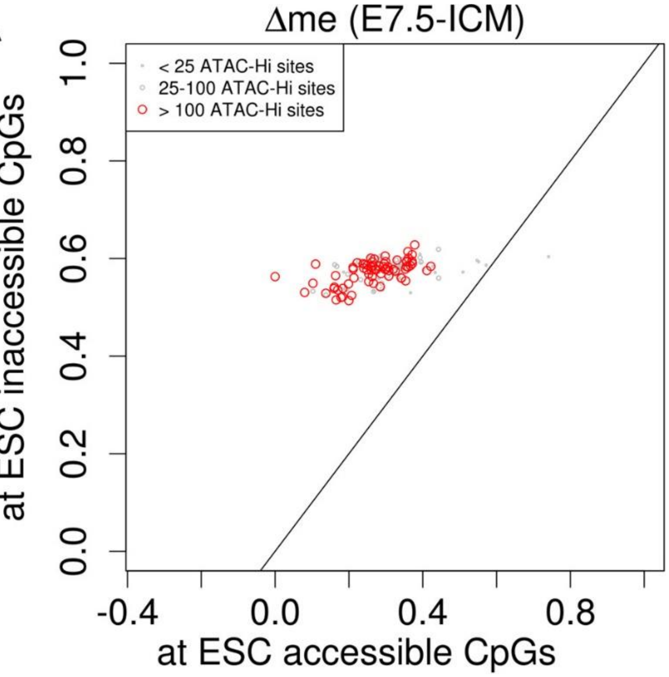

Δme (E7.5-ICM)
< 25 ATAC-Hi sites
25-100 ATAC-Hi sites
> 100 ATAC-Hi sites
at ESC inaccessible CpGs
at ESC accessible CpGs
0.0
0.2
0.4
0.6
0.8
1.0
-0.4
0.0
0.4
0.8


B

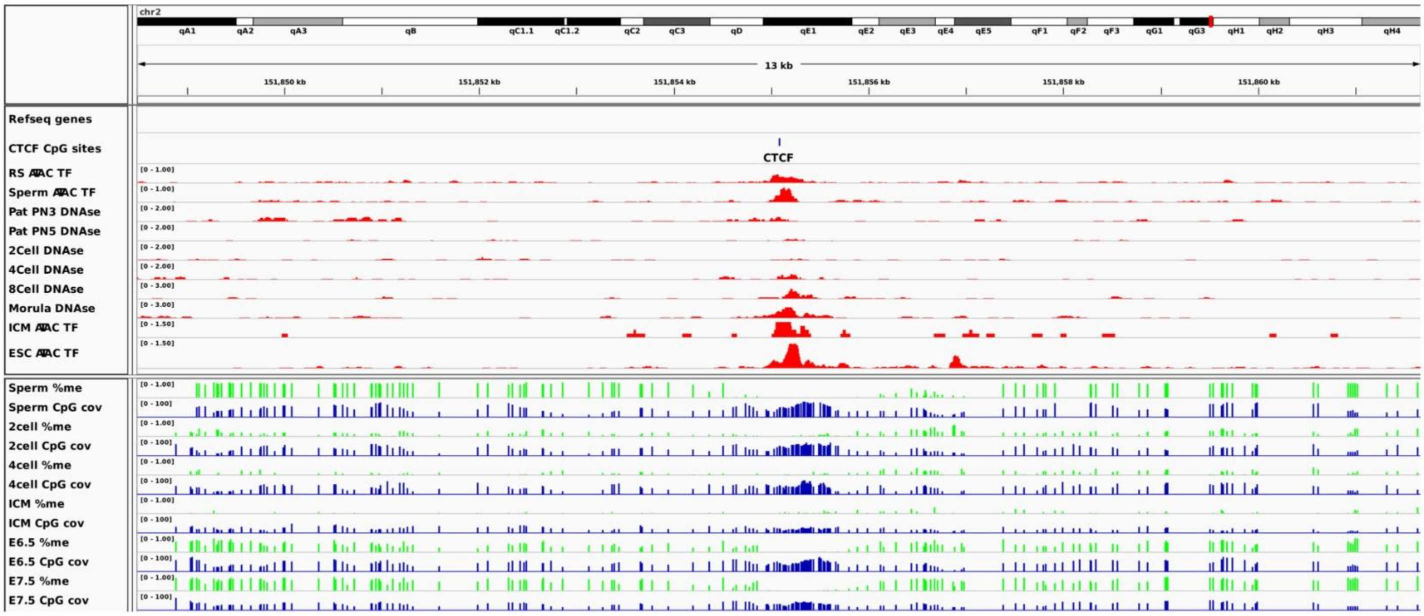

chr2
13 kb
151,850 kb
151,852 kb
151,854 kb
151,856 kb
151,858 kb
151,860 kb
Refseq genes
CTCF CpG sites
CTCF
RS ATAC TF
Sperm ATAC TF
Pat PN3 DNAse
Pat PN5 DNAse
2Cell DNAse
4Cell DNAse
8Cell DNAse
Morula DNAse
ICM ATAC TF
ESC ATAC TF
Sperm %me
Sperm CpG cov
2cell %me
2cell CpG cov
4cell %me
4cell CpG cov
ICM %me
ICM CpG cov
E6.5 %me
E6.5 CpG cov
E7.5 %me
E7.5 CpG cov


C

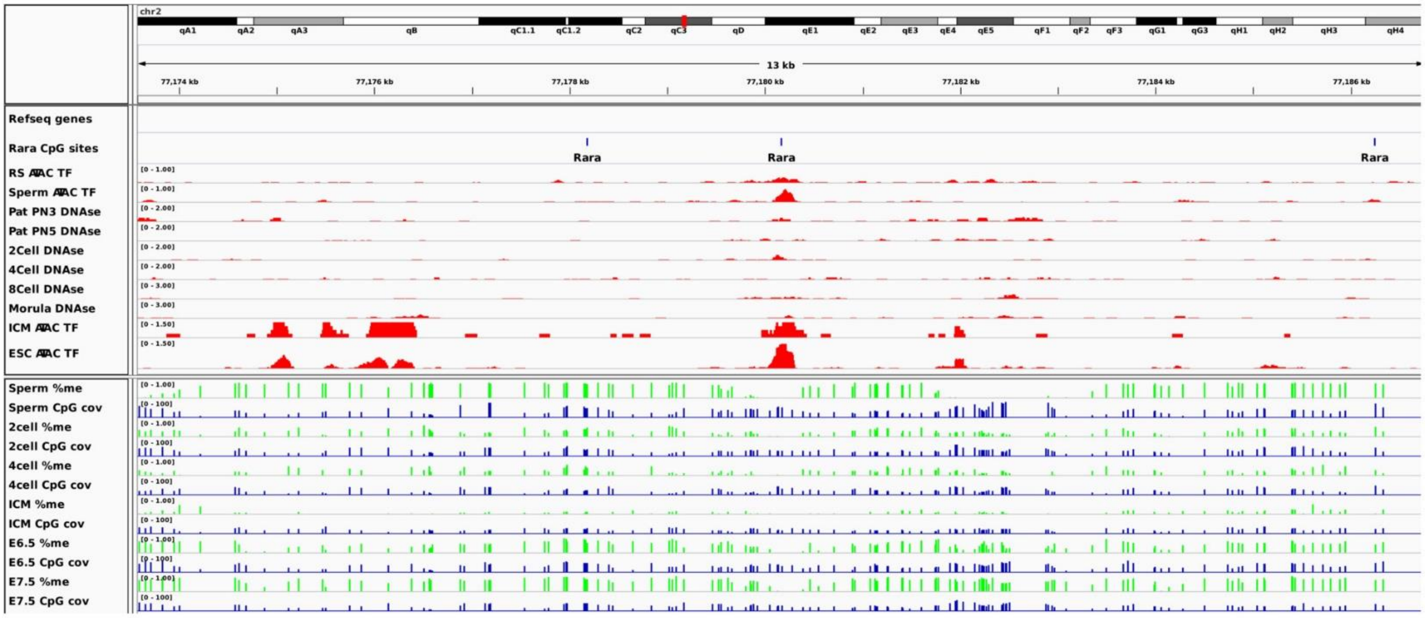

chr2
13 kb
77,174 kb
77,176 kb
77,178 kb
77,180 kb
77,182 kb
77,184 kb
77,186 kb
Refseq genes
Rara CpG sites
Rara
Rara
Rara
RS ATAC TF
Sperm ATAC TF
Pat PN3 DNAse
Pat PN5 DNAse
2Cell DNAse
4Cell DNAse
8Cell DNAse
Morula DNAse
ICM ATAC TF
ESC ATAC TF
Sperm %me
Sperm CpG cov
2cell %me
2cell CpG cov
4cell %me
4cell CpG cov
ICM %me
ICM CpG cov
E6.5 %me
E6.5 CpG cov
E7.5 %me
E7.5 CpG cov

**Figure S2.** Related to Figure 2. **A** Scatterplot of changes in DNA methylation between E14.5m and E16.5m PGCs. Each point represents the average methylation change (fraction meCpG in E16.5m minus E13.5m PGCs) at CpGs overlapping a binding sequence for a specific TF. The x-axis gives the average value for E14.5m PGC DNase-Hi sites, while the y-axis gives the average value for DNase-Lo sites. **B,C** Example regions showing TF accessibility and DNA methylation in preimplantation embryos at E14.5m PGC DNAse-Hi sites. CpG cov indicates the BS-seq read coverage. TF binding sequences called by fimo that overlap the peak summit in E14.5m are shown.

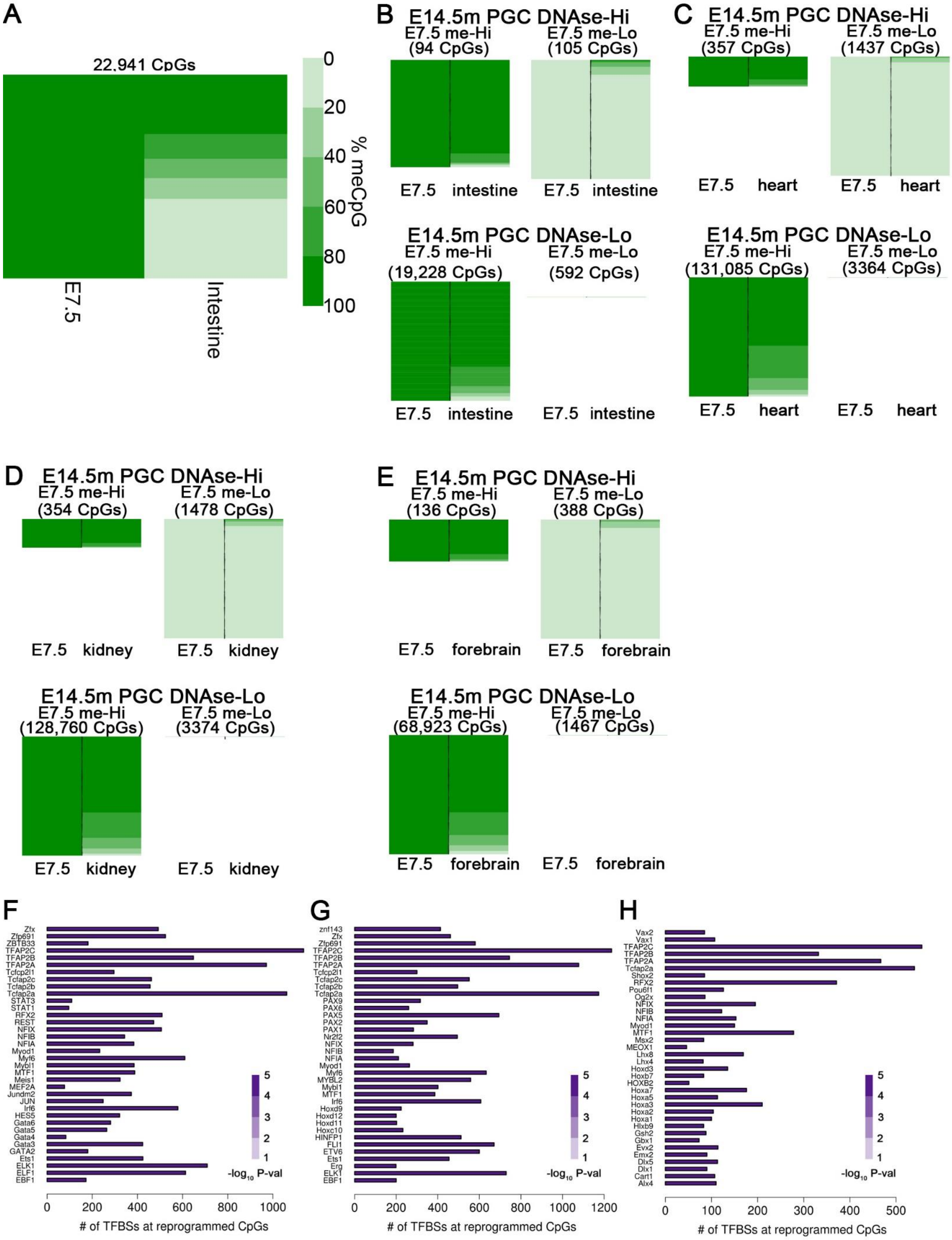

A
22,941 CpGs
E7.5
Intestine
0
20
40
60
80
100
% meCpG
B
E14.5m PGC DNAse-Hi
E7.5 me-Hi
(94 CpGs)
E7.5 me-Lo
(105 CpGs)
E7.5 intestine
E7.5 intestine
E14.5m PGC DNAse-Lo
E7.5 me-Hi
(19,228 CpGs)
E7.5 me-Lo
(592 CpGs)
E7.5 intestine
E7.5 intestine
C
E14.5m PGC DNAse-Hi
E7.5 me-Hi
(357 CpGs)
E7.5 me-Lo
(1437 CpGs)
E7.5 heart
E7.5 heart
E14.5m PGC DNAse-Lo
E7.5 me-Hi
(131,085 CpGs)
E7.5 me-Lo
(3364 CpGs)
E7.5 heart
E7.5 heart
D
E14.5m PGC DNAse-Hi
E7.5 me-Hi
(354 CpGs)
E7.5 me-Lo
(1478 CpGs)
E7.5 kidney
E7.5 kidney
E14.5m PGC DNAse-Lo
E7.5 me-Hi
(128,760 CpGs)
E7.5 me-Lo
(3374 CpGs)
E7.5 kidney
E7.5 kidney
E
E14.5m PGC DNAse-Hi
E7.5 me-Hi
(136 CpGs)
E7.5 me-Lo
(388 CpGs)
E7.5 forebrain
E7.5 forebrain
E14.5m PGC DNAse-Lo
E7.5 me-Hi
(68,923 CpGs)
E7.5 me-Lo
(1467 CpGs)
E7.5 forebrain
E7.5 forebrain
F
Zfx
Zfp691
ZBTB33
TFAP2C
TFAP2B
TFAP2A
Tcfcp2l1
Tcfap2c
Tcfap2b
Tcfap2a
STAT3
STAT1
RFX2
REST
NFIX
NFIB
NFIA
Myod1
Myf6
Mybl1
MTF1
Meis1
MEF2A
Jundm2
JUN
Irf6
HES5
Gata6
Gata5
Gata4
Gata3
GATA2
Ets1
ELK1
ELF1
EBF1
0 200 400 600 800 1000
### of TFBSs at reprogrammed CpGs
5 4 3 2 1
$-log_{10}$ P-val
G
znf143
Zfx
Zfp691
TFAP2C
TFAP2B
TFAP2A
Tcfcp2l1
Tcfap2c
Tcfap2b
Tcfap2a
PAX9
PAX6
PAX5
PAX2
PAX1
Nr2f2
NFIX
NFIB
NFIA
Myod1
Myf6
MYBL2
Mybl1
MTF1
Irf6
Hoxd9
Hoxd12
Hoxd11
Hoxc10
HINFP1
FLI1
ETV6
Ets1
Erg
ELK1
EBF1
0 200 400 600 800 1000 1200
### of TFBSs at reprogrammed CpGs
5 4 3 2 1
$-log_{10}$ P-val
H
Vax2
Vax1
TFAP2C
TFAP2B
TFAP2A
Tcfap2a
Shox2
RFX2
Pou6f1
Og2x
NFIX
NFIB
NFIA
Myod1
MTF1
Msx2
MEOX1
Lhx8
Lhx4
Hoxd3
Hoxb7
HOXB2
Hoxa7
Hoxa5
Hoxa3
Hoxa2
Hoxa1
Hlxb9
Gsh2
Gbx1
Evx2
Emx2
Dlx5
Dlx1
Cart1
Alx4
0 100 200 300 400 500
### of TFBSs at reprogrammed CpGs
5 4 3 2 1
$-log_{10}$ P-val

**Figure S3.** Related to Figure 4. **A** Heatmap of CpG methylation percentage for all CpGs with > 80% methylation in E7.5 embryos that overlap with a published ATAC-seq peak in either embryonic, fetal, or adult intestine. The legend in A also applies to panels B-E. **B** Heatmap as described in Figure 4A, except that only CpGs that do not overlap a published ATAC-seq peak from fetal, embryonic, or adult intestinal tissue are considered. **C-E** Heatmaps as in Figure 4A, except that instead of adult intestine, methylation is examined from BS-seq of neonatal heart (C), kidney (D), and forebrain (E). **F-H** Bar plots showing the number of motifs and the statistical significance of motif enrichment for the indicated TFs at CpGs with > 80% methylation in E7.5 embryos and < 20% methylation in neonatal heart (F), kidney (G), and forebrain (H).

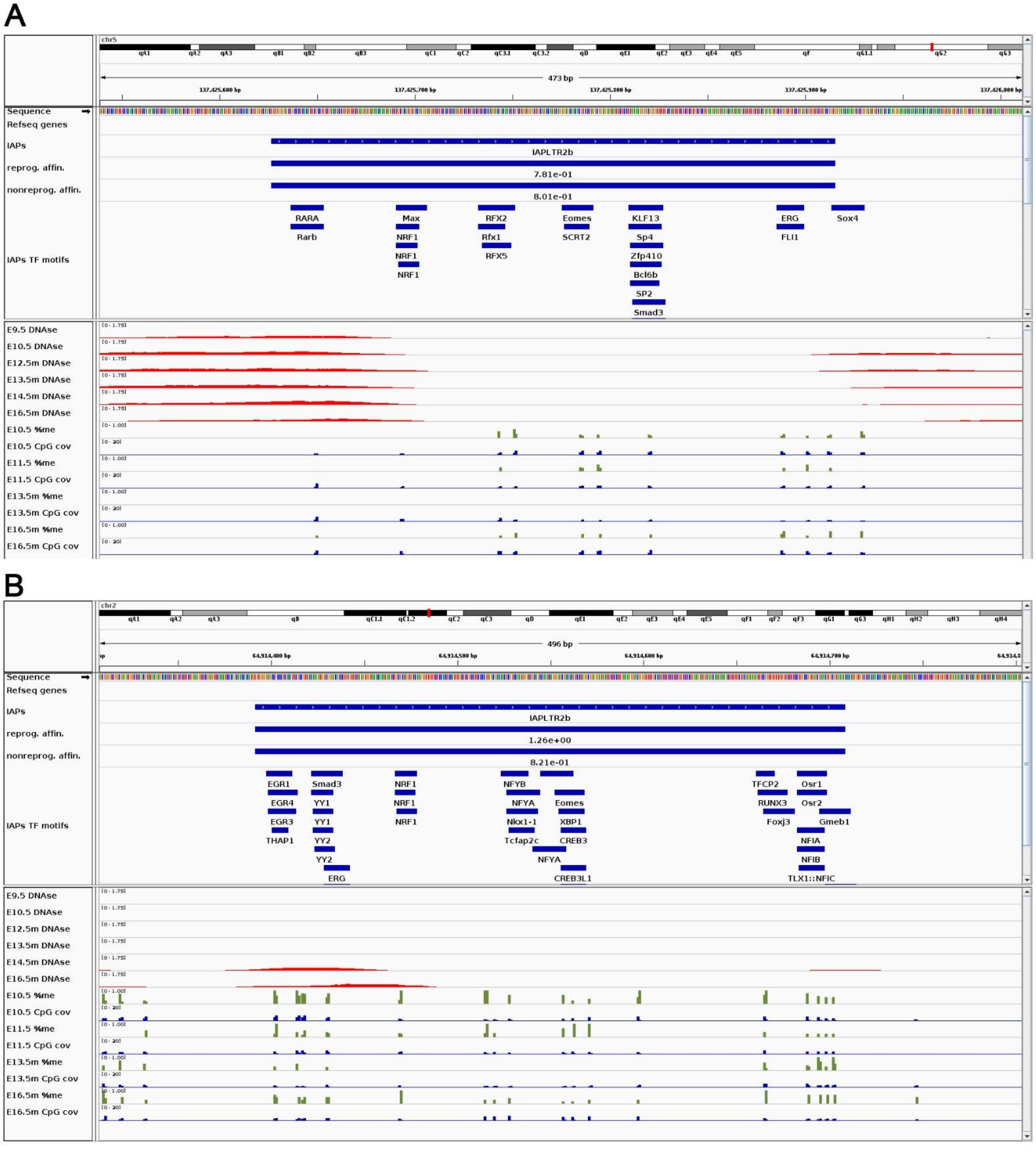


**Figure S4.** Related to Figure 6. **A,B** Examples showing DNase-seq and DNA methylation levels during PGC development at IAP LTRs that have evidence of TF binding. Tracks showing the highest TRAP affinity of the E16.5m PGC putative reprogramming TFs, and of E16.5m PGC non-reprogramming TFs, at the whole IAP, are displayed under the IAP track.
